# MAST: Phylogenetic Inference with Mixtures Across Sites and Trees

**DOI:** 10.1101/2022.10.06.511210

**Authors:** Thomas KF Wong, Caitlin Cherryh, Allen G Rodrigo, Matthew W Hahn, Bui Quang Minh, Robert Lanfear

## Abstract

Hundreds or thousands of loci are now routinely used in modern phylogenomic studies. Concatenation approaches to tree inference assume that there is a single topology for the entire dataset, but different loci may have different evolutionary histories due to incomplete lineage sorting, introgression, and/or horizontal gene transfer; even single loci may not be treelike due to recombination. To overcome this shortcoming, we introduce the mixture across sites and trees (MAST) model, which uses a mixture of bifurcating trees to represent multiple histories in a single concatenated alignment. The MAST model allows each tree to have its own topology, branch lengths, substitution model, nucleotide or amino acid frequencies, and model of rate heterogeneity across sites. We implemented the MAST model in a maximum-likelihood framework in the popular phylogenetic software, IQ-TREE. Simulations show that we can accurately recover the true model parameters, including branch lengths and tree weights (i.e. frequencies) for a given set of tree topologies. We also show that we can use standard statistical inference approaches to reject a single-tree model when data are simulated under multiple trees (and vice versa). We applied the MAST model to multiple primate datasets and found that it can recover the signal of incomplete lineage sorting in the Great Apes, as well as the asymmetry in minor trees caused by introgression among several macaque species. When applied to a dataset of four Platyrrhine species for which standard concatenated maximum likelihood and gene tree approaches disagree, we find that MAST gives the highest weight to the tree favored by gene tree approaches. These results suggest that the MAST model is able to analyse a concatenated alignment using maximum likelihood, while avoiding some of the biases that come with assuming there is only a single tree. The MAST model can therefore offer unique biological insights when applied to datasets with multiple evolutionary histories. We discuss how it can be extended in the future.

## Introduction

Molecular phylogenetics aims to infer phylogenetic trees, often from aligned DNA or amino acid (AA) sequencing data. Many popular phylogenetic tools are designed to infer a single tree from a multiple sequence alignment, using one of a number of approaches such as maximum likelihood (e.g. RAxML (1), IQ-TREE (2), PhyML (3)), Bayesian inference (e.g. MrBayes (4), BEAST (5)), maximum parsimony (e.g. MPBoot (6), matOptimize (7), TNT (8)), or distance methods (e.g. BioNJ (9), FastME (10), QuickTree (11), RapidNJ (12)). The assumption that the data can be represented as a single tree is appropriate when analysing a single non-recombining locus. However, there are many situations where this “treelikeness” assumption is violated. For example, an alignment of a single locus may contain one or more recombination events in its history, such that different regions of the alignment follow different trees. More generally, it is well known that different genomic loci may have evolved under different trees due to biological processes including incomplete lineage sorting (ILS), hybridisation/introgression, and horizontal gene transfer (13, 14). Since modern phylogenomic datasets now routinely contain hundreds or thousands of loci, a great deal of work has focused on developing methods and software that relax the treelikeness assumption (15). Most existing approaches that account for complex histories in large datasets focus on inferring either species trees or species networks, either from a single concatenated alignment or from many individual locus alignments or individual locus trees. Many of the most popular approaches for inferring species trees are based on the multi-species coalescent model (MSC) or are consistent with the MSC, and can infer a species tree while accounting for ILS among loci (e.g. SNAPP (16), ASTRAL-III (17), MP-EST (18), SVD-Quartets (19), *BEAST (20), *BEAST2 (21)). More recent work has extended the MSC to account for a broader range of processes that can cause reticulations in the underlying species tree. These methods use models referred to as the multi-species network coalescent (or MSNC), and typically infer a species network that represents both the vertical inheritance and horizontal exchange of genetic material among evolving lineages (e.g. PhyloNet (22), PhyloNetworks (23), SpeciesNetwork (24), and BPP (25)).

In this study, we present a different solution to the problem of accounting for multiple histories in a single sequence alignment: the mixtures across sites and trees (MAST) model. The MAST model is an example of a *multitree* mixture model (26), because it uses mixtures of bifurcating trees to represent the multiple histories present in a dataset. In phylogenetic mixture models, a number of sub-models (known as classes) are estimated from the data and the likelihood of each site in the alignment is calculated as the weighted sum of the likelihood for that site under each sub-model (Figure 1). Mixture models have been widely used in phylogenetic inference, including in rate heterogeneity across site models (2, 27), profile mixture models (e.g. the CAT model (28)), mixtures of substitution rate matrices (e.g. the LG4M and LG4X models (29)), and mixtures of branch lengths (e.g. the GHOST model (30)). The MAST model extends the use of mixture models to tree topologies. It is best seen as a generalisation of a standard concatenated phylogenetic analysis. In a standard concatenated phylogenetic analysis, we assume that the history of the entire alignment is represented by a single bifurcating phylogenetic tree (i.e. we make the treelikeness assumption). With the MAST model we relax this assumption and represent the history of the alignment with a mixture of any number of trees. Given an alignment and a collection of tree topologies that contain the same tip labels as that alignment, the MAST model estimates the likelihood of each site under each tree, the maximum-likelihood weights of each of the input trees, the branch lengths of the trees, and the other free parameters of the substitution model. In this way, it has many of the advantages over concatenation approaches, but can still accommodate underlying discordance in the alignment (31).

**Fig. 1.**
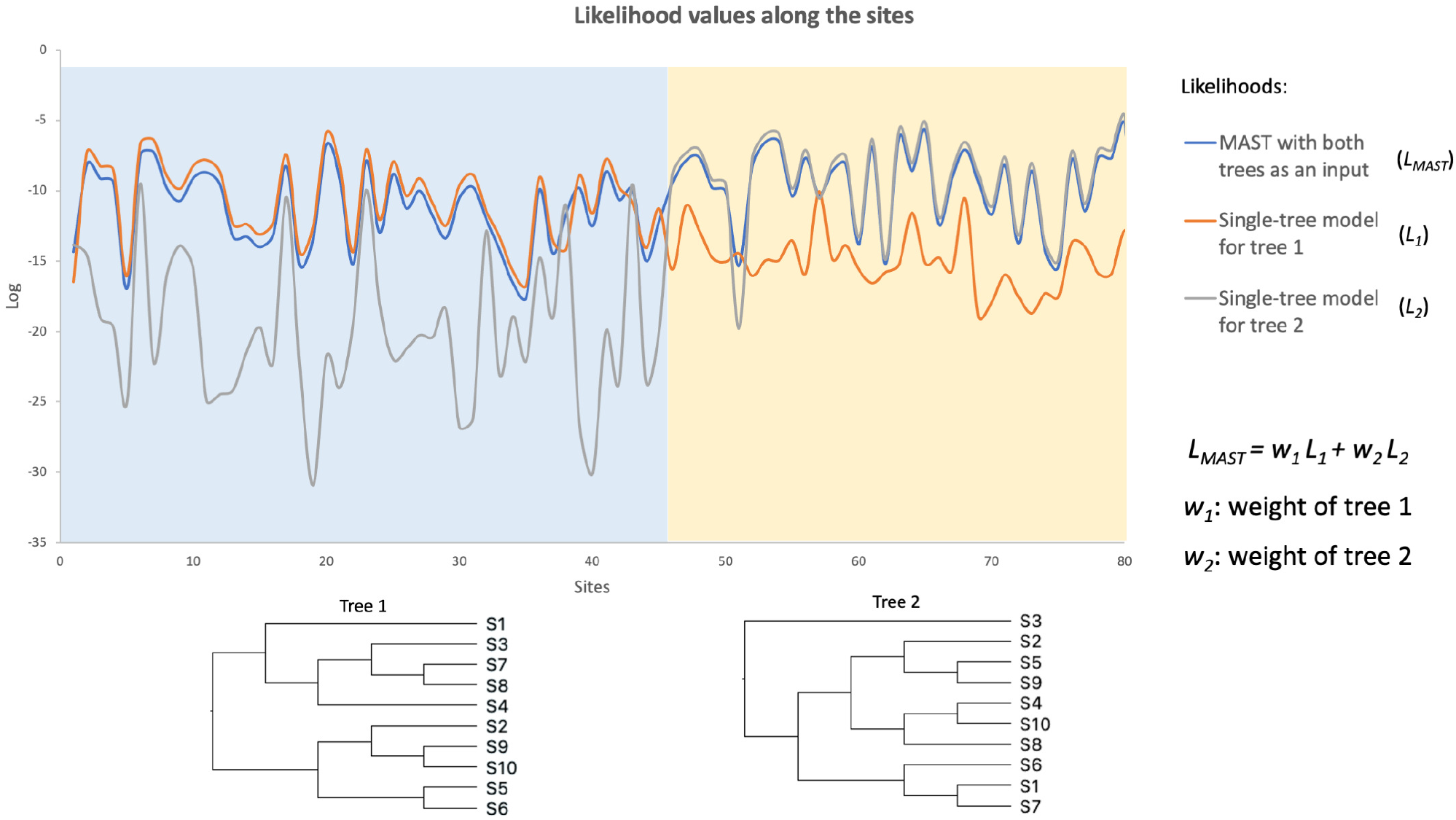
An example illustrating the MAST model. Two regions (of length 45 bp and 35 bp) were simulated under two different topologies, each with ten taxa. The curves at the top show the site likelihoods (on a log scale) computed under tree 1 (*L*_1_), tree 2 (*L*_2_), and the MAST model (*L_MAST_*). *L_MAST_* is calculated as the weighted sum of *L*_1_ and *L*_2_, where the weight parameters *w*_1_ and *w*_2_ will be estimated by the MAST model. This toy example shows that the *L_MAST_* curve matches the *L*_1_ curve for region 1 and the *L*_2_ curve for region 2 with high site likelihoods, demonstrating the ability of the MAST model to predict the true underlying evolution of this data.

The MAST model differs from species tree and species network models in a number of ways. First, as opposed to many MSC and MSNC approaches, the MAST model does not explicitly model biological processes such as ILS, introgression, or horizontal gene transfer. Instead, the MAST model is process-agnostic and simply seeks to calculate the relative weights of multiple histories from the input data. This is a limitation in the sense that the output of the MAST model does not contain direct estimates of many evolutionary parameters of interest, such as the number of hybridisation events, their location on the species tree, or ancestral population sizes. On the other hand, this difference may be seen as a strength because the MAST model can represent all biological processes or bioinformatic errors (such as the accidental inclusion of paralogs) which can cause the treelikeness assumption to be violated for a given alignment. Second, the MAST model differs from previous approaches because it calculates the likelihood of every site under every tree in the mixture, and thus allows fine-grained post-hoc analysis of the data. For example, while a species network can estimate the proportion of the genome that may have been involved in an introgression event, the MAST model represents the weights of the relevant tree topologies as well as the association between the sites of the alignment and the event.

In this paper, we first describe the mathematical basis of the MAST model and its implementation in IQ-TREE. This implementation allows us to estimate model parameters and branch lengths for a given set of fixed tree topologies. We then perform extensive simulations to evaluate the accuracy and the limitations of the MAST model. Finally, we demonstrate the use of the MAST model on four empirical datasets of primates to show that it recapitulates results from well-studied clades. We also highlight the advantages of MAST over standard phylogenetic analysis methods when applied to these datasets.

## Materials and Methods

### The MAST model

In a standard concatenation analysis (such as that performed by IQ-TREE (32) or RAxML (1)), it is assumed that every site in the concatenated alignment comes from a single phylogenetic tree, which consists of a topology and branch lengths. In this framework, maximum likelihood (ML) approaches seek to find the model of sequence evolution, tree topology, and branch lengths that maximize the likelihood of the observed alignment. The MAST model generalizes this framework by assuming that each site in the alignment comes from a mixture of *m* trees. Each tree has its own topology and branch lengths, and the trees may have independent or shared substitution models (e.g. the general time reversible (GTR) model (33)), a set of nucleotide or amino-acid frequencies, and a rate heterogeneity across sites (RHAS) model (e.g. the +G or +I+G models). In what follows we first describe the case in which each tree has an independent substitution model, set of nucleotide or amino acid frequencies, and RHAS model.

### Model description

The MAST model consists of *m* classes where each class comprises a bifurcating tree topology *T_j_*. For the *j*-th class, *λ_j_* is defined as the set of branch lengths on *T_j_, R_j_* as the relative substitution rate parameters, *F_j_* as the set of nucleotide or amino-acid frequencies, *H_j_* as the rate heterogeneity model, and *w_j_* as the class weight (*w_j_* > 0, 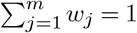). Given a multiple sequence alignment, *A*, we define *L_ij_* as the likelihood of the data observed at *i*-th site in *A* under the *j*-th class of the MAST model. *L_ij_* can be computed using Felsenstein’s pruning algorithm (34). The likelihood of the *i*-th site, *L_i_*, is the weighted sum of the *L_ij_* over the *m* classes:

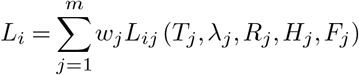

The full log-likelihood *l* over all *N* alignment sites, which are assumed to be independent and identically distributed (iid), is:

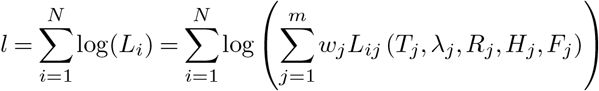

This formula is very similar to the formulation of the GHOST model (30), which allows for mixtures of branch lengths on a single topology, and differs only insofar as the final sum here is across the *m* tree topologies and their associated branch lengths, versus the *m* sets of branch lengths on a single topology in the GHOST model.

In the implementation of the MAST model we describe here we assume that we know the topologies of all of the *m* trees ahead of time, for example, the set of gene tree topologies observed among the genomes, or the set of possible trees that should appear under the MSC model. We then estimate the relative weights (i.e. proportions) of each topology, optimize the branch lengths of each topology, the parameters of the evolutionary model, and the nucleotide or amino-acid frequencies for each tree. We consider extensions of the model when the tree topologies are not given in the Discussion.

### Linked and unlinked MAST submodels

In standard phylogenetic analyses we estimate a single tree with an associated set of branch lengths, along with the parameters of the substitution model, the base or amino acid frequencies, and the rate heterogeneity across sites (RHAS) model. In the most general MAST model introduced above (submodel 1 in Figure 2), the tree, the branch lengths of that tree, the substitution model, the base or amino acid frequencies, and the RHAS model can all vary in each class, and the weight of that class pertains to the full set of free parameters associated with that class. We say that all parameters are unlinked across classes in this model. We also allow for five more-restrictive models in which the parameters of the substitution models, the vectors of base or amino acid frequencies, or the RHAS model can be linked across all *m* classes of trees and their associated branch lengths. The most restricted model (submodel 6 in Figure 2) links the parameters of all three of these components of the model across all *m* classes of trees and their associated branch lengths. In this model, the estimated weights therefore pertain only to the trees and their branch lengths in each of the *m* classes, because these are the only parameters allowed to differ among classes. This framework allows for the comparison of models with likelihood ratio tests or other information criteria (35).

**Fig. 2.**
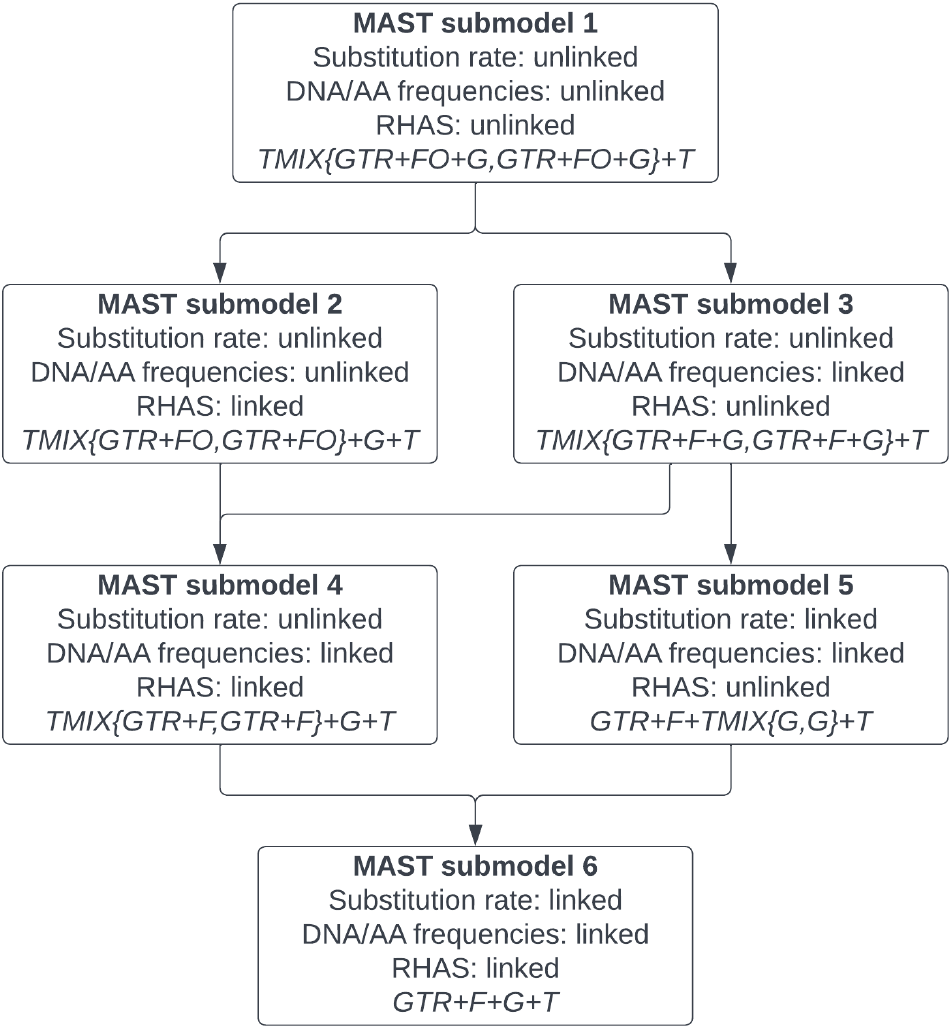
A hierarchy of six MAST submodels currently implemented in IQ-TREE. The term ‘unlinked’ means the parameters can differ across mixture classes, while ‘linked’ means the parameters are restricted to be equal across all classes. The last line in each box shows the name of the model that can be used directly as input in IQ-TREE via -m option, assuming two classes with a GTR substitution model and Gamma RHAS model for each class. The arrows indicate the nestedness between the submodels; for example, submodel 4 is nested within both submodels 2 and 3, while submodel 6 is nested within both submodels 4 and 5. Note that two submodels are missing (i.e. substitution rate: linked; DNA/AA frequencies: unlinked; RHAS: linked/unlinked) due to a non-trivial implementation.

### Model parameter estimation for fixed topologies

Given a set of fixed topologies, *T*_1_,…, *T_m_*, the challenge is to optimize all of the parameters without getting stuck in local optima. We employ both the expectation-maximization (EM) algorithm (36) and the Broyden-Fletcher-Goldfarb-Shanno (BFGS) algorithm (37) to estimate the MAST model parameters. Taking advantage of the existing parameter optimization algorithms implemented in IQ-TREE, our workflow (Figure 3) operates as follows. To begin, for class *j*, the substitution model *R_j_* and the nucleotide or amino-acid frequencies *F_j_* are initialized as a Jukes-Cantor (JC) model (i.e. 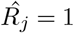 and uniform frequencies *F_j_*), and the branch lengths *λ_j_* are initialized as the maximum parsimony (Fitch 1971) branch lengths of the tree *T_j_*. To obtain some sensible initial values of the tree weights, we first compute the parsimony scores for each tree topology along all the sites. For each of the sites with different parsimony scores between the tree topologies, we then check which tree topology has the minimum parsimony score and assign the site to that tree. The tree weights are then initialized according to the proportion of the sites assigned to each of the trees. If all sites have the same parsimony scores across all the trees, then the tree weights are initialized to be equal.

**Fig. 3.**
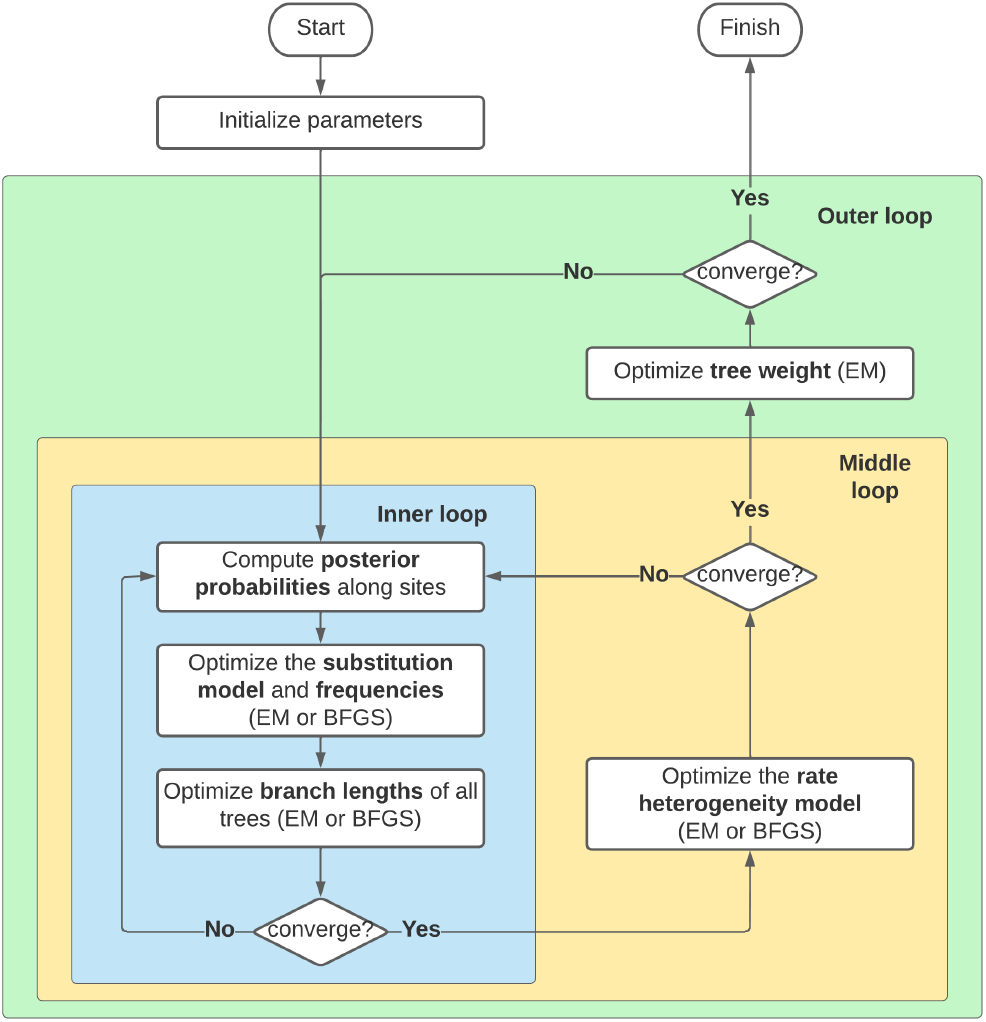
Optimization flow chart for the MAST model in IQ-TREE. The optimization workflow includes an outer loop, a middle loop, and an inner loop of iterations. The inner loop optimizes the substitution model, nucleotide frequencies, and branch length of the trees; the middle loop optimizes the rate heterogeneity model; the outer loop optimizes the tree weights. The EM algorithm is used to optimize the individual unlinked parameters of each tree and the BFGS algorithm is used to optimize the linked parameters. The iterations continue until the likelihood value converges.

Having established the starting values for all the parameters in the model, we then optimize them. The optimization of each class of model parameters is done sequentially. Figure 3 summarizes the workflow of the optimization. Our optimization workflow includes an outer loop, a middle loop, and an inner loop of iterations. The inner loop optimizes the substitution model, nucleotide frequencies, and branch length of the trees; the middle loop optimizes the rate heterogeneity model; the outer loop optimizes the tree weights. This optimisation continues to iterate until the resulting loglikelihood value converges, where convergence is defined as the increment of the log-likelihood value in the current iteration falling below some threshold *ϵ* (which we set to 0.0001). To optimize the unlinked parameters of each tree in the mixture model, we use an EM algorithm similar to that used in the GHOST model (30).

In detail, our calculations are as follows. Define *p_i,j_* as the posterior probability of site *D_i_* evolving under a tree *T_j_*. The value of *p_i,j_* is computed by the following equation:

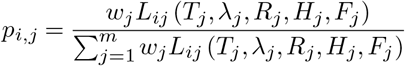

The expectation of the log-likelihood value (*l_j_*) of tree *j* over all the sites:

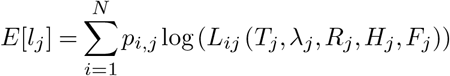

In every iteration, by fixing the posterior probabilities *p_i,j_*, we optimize the tree weights, the branch lengths, the unlinked substitution rate models, and the unlinked rate heterogeneity models of all trees one-by-one to maximize the expected likelihood value. The tree weights are then updated by averaging the probabilities over all the *N* sites. That is, the new weight of class *j* is the mean posterior probability of each site belonging to class *j*:

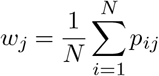

For the linked models (submodels 2-6 in Figure 2) the EM algorithm cannot be applied to the optimisation of the linked parameters shared between the classes. Thus, we optimize the parameters of the linked substitution rate model *R*, the linked nucleotide or amino acid frequencies *F*, and the linked rate heterogeneity model *H* using the BFGS algorithm in IQ-TREE.

### Simulations

Having implemented the MAST model in IQ-TREE, we next used simulated data to test the performance of the MAST model under a wide range of scenarios. The first and second simulation experiments test the accuracy of the unlinked and linked MAST models when the true model is specified. The third simulation experiment simulates data with varying levels of introgression to compare the performance of standard (i.e. single-tree) concatenation methods to the performance of the MAST model. The fourth simulation experiment examines the performance of the MAST model when an incorrect model is specified, by applying an unlinked MAST model with different numbers of trees to an alignment simulated under a single tree.

### Simulations 1 & 2: Parameter estimation under the true model for unlinked and linked MAST model (submodel 1 & submodel 6)

These simulations are designed to ask whether our implementation of the MAST model in IQ-TREE is capable of estimating accurate tree weights, branch lengths, and other model parameters when the model used for inference matches the model used for simulation. We simulated alignments under the completely unlinked MAST model (submodel 1 in Figure 2; simulation 1) and the completely linked MAST model (submodel 6 in Figure 2; simulation 2), and provided IQ-TREE with the set of true tree topologies from the mixture, as well as the true model of molecular evolution (e.g. GTR+G), and the correct MAST model (i.e. submodel 1 or 6). We then measured the accuracy of our implementation by recording the estimated tree weights, branch lengths, substitution model parameters, and nucleotide frequencies, and comparing them to the values used to simulate the data.

We simulated alignments from mixtures of *m* of trees with different numbers (*t*) of taxa, where *m* ∈ {1,2,3,5,10} and *t* ∈ {6,7,10,20}. We performed 100 replicate simulations for every combination of *m* and *t*, for a total of 2000 simulated datasets per experiment. For each simulated dataset, the *m* random trees with *t* tips were simulated by IQ-TREE version 2 (2) with the option ‘-r’. This simulates a random Yule tree with branch lengths drawn from an exponential distribution with a mean of 0.1, a minimum of 0.001, and a maximum of 0.999.

The parameters of the GTR model *R* were the values of six random integers uniformly drawn between 5 and 50 and normalized according to the value of G↔T such that the G↔T rate was always equal to 1.0. The gamma rate *H* was drawn randomly from an exponential distribution with a mean of 1.0. The set of nucleotide frequencies *F* was the proportions between four integers uniformly drawn between 1 and 10. Different *R, H*, and *F* were simulated over the trees in the first simulation experiment, while the same *R*, *H*, and *F* were shared among the trees in the second simulation experiment. The alignments were then simulated according to the tree, the GTR model, and the gamma rate using AliSim (38). Each simulated dataset contained 100k bases, regardless of the number of trees *m*. The proportions of the lengths of each of the *m* alignments simulated from each of the *m* trees were the ratios of *m* random integers drawn from a uniform distribution between 1 and 10.

To assess the accuracy of the parameter estimates, we calculated the root-mean-squared error (RMSE) of each estimated parameter when compared to its value in the simulation. For each dataset, we compared the statistical fit of the MAST model to that of a standard single-tree model by comparing the BIC value (*BIC*) of the MAST model to the BIC value(*BIC*0) of a standard single-tree model.

### Simulation 3: Introgression

To examine the performance of the MAST model in a biologically motivated setting, we simulated alignments on 4-taxon trees with different levels of introgression and then used both a single-tree model and the linked MAST model (i.e. submodel 6) to analyse them. Each dataset was simulated from a rooted 4-taxon tree shown in Supplementary Figure S4A. Using this tree we simulated 1500 gene trees with introgression rate *r* from lineage 2 to lineage 4 (Supplementary Figure S4A) using the program *ms* (39), where *r* ∈ {0.0,0.1,0.2,…, 0.9,1.0}. When the introgression rate is zero, the largest fraction of the gene trees will match the species tree *T*_*E*1_ and the frequency of the two minor trees, *T*_*E*2_ and *T*_*E*3_, are expected to be equal. As the introgression rate increases, the frequency of the tree matching the introgression history, *T*_*E*2_, will increase, and the frequency of the other two trees will decrease. The MAST model should reflect these patterns in the tree weights calculated from a concatenated alignment of all 1500 genes, without the need to know the boundaries between the individual loci. The benefit of this approach when applied to an empirical dataset is that it overcomes concerns about ‘concatalesence’, in which unaccounted-for recombination within loci can bias estimates of gene tree frequency calculated by building trees for each locus (40). Since ms uses a coalescent model, we rescaled the branch lengths from coalescent units to units appropriate for simulating alignments (i.e. substitutions per site) by multiplying all branch lengths by 0.002, selected to result in branch lengths similar to those recovered from our analyses of empirical dataset 4 (see below). For each simulated gene tree, we used AliSim (38) to simulate a 1000bp alignment using the GTR+G model with parameters equal to those reported by IQ-TREE for our analysis of empirical dataset 4 (see below). Concatenating all the single-locus alignments resulted in an alignment of 1,500,000bp. We performed 100 replicate simulations at every *r*, for a total of 1100 simulated datasets. We then applied the linked MAST model (submodel 6 in Figure 2) to these data, with the input trees comprised of all three possible unrooted trees of the four taxa in Supplementary Figure S4B.

### Simulation 4: Parameter estimation under misspecified models

We next sought to examine the performance of the MAST model when the underlying data were simulated under a single tree *T*, but the data were analysed under a MAST model with *m* > 1 i.e. a misspecified model with more than one tree. To do this, we simulated data under a single tree topology, and then applied MAST submodel 6 (Figure 2) where the *m* trees included the true tree *T* and also *m* – 1 additional tree topologies that differed from *T*. This simulation is designed to examine the case where a researcher includes the primary tree in a MAST model (e.g. a tree derived from a single-tree concatenated ML analysis, or an MSC analysis) but additionally includes some hypothesized trees in the model that have no support in the underlying data.

Each simulated dataset comprised 20k base pairs and was simulated on a single tree with different numbers (*t*) of taxa, where *t* ∈ {6,7,10,20}; other simulation parameters were as above for MAST model 6. We performed 100 replicate simulations at every *t*, for a total of 400 simulated datasets. We then analysed each simulated dataset under MAST submodel 6. To simulate each of the additional *m* – 1 tree topologies in each MAST model, we sequentially performed *k* random subtree pruning and regrafting (SPR) moves on the true tree *T*. The completely linked MAST model was then applied by inputting the actual tree topology as well as the other *m* – 1 different tree topologies that all are *k*-SPR moves from that tree, where *m* ∈ {2,3,5,10} and *k* ∈ {1,2,3}. Analysing each of the 400 simulated datasets under 12 combinations of *m* and *k* gives a total of 4800 analyses.

To evaluate the performance, among the 100 replicates, we recorded how many times the true topology had the maximum tree weight. As above we also compared the BIC value (*BIC*) reported by the MAST model with the BIC value (*BIC*0) under the true model, i.e. when the dataset was analysed under the single true tree *T*.

### Applications to empirical data

In addition to testing the MAST model on simulated data, we also applied it to four empirical datasets (Table 1). All of these datasets are subsets of a single dataset comprising 1730 single-gene alignments from 26 primates (41). The first two empirical datasets we used are simple four-taxon datasets, in which it is trivial to supply the MAST model with all three possible unrooted trees, and for which the expected tree weights have been estimated in previous research. In the other two empirical experiments, a standard single-tree model was first used to infer a topology for every gene in the dataset. Then, the set (or subset) of most commonly inferred gene trees were used as the set of input topologies for the MAST model when analysing a concatenated alignment of all the single-gene alignments. In order to find out whether the MAST model has a better fit to the data compared with the standard single-tree model, we analysed multiple different submodels of MAST (Figure 2). We compared the lowest BIC values from these models to the BIC value calculated using the standard single-tree model on the same alignments.

**Table 1.** The fourempirical datasets analysed here

| Empirical datasets | Species | # of genes | Total length |
| --- | --- | --- | --- |
| A | *Homo sapiens*, *Pan troglodytes*,<br>*Gorilla gorilla*, *Pongo abelii* | 1,595 | 1,618,506 |
| B | *Macaca fascicularis*, *Macaca mulatta*,<br>*Macaca nemestrina*, *Colobus angolensis palliatus* | 1,599 | 1,629,163 |
| C | *Homo sapiens*, *Pan troglodytes*, *Gorilla gorilla*,<br>*Macaca fascicularis*, *Macaca mulatta*, *Macaca nemestrina* | 1,556 | 1,576,852 |
| D | *Callithrix jacchus*, *Aotus nancymae*, *Saimiri boliviensis*,<br>*Cebus capucinus imitator*, *Macaca mulatta* | 1,557 | 1,610,755 |

The first dataset (‘A’) includes the well-studied four-taxon grouping of human, chimpanzee, gorilla, and orangutan. Previous studies have shown that all three possible unrooted gene trees of four taxa (Figure 4; orangutan is considered an outgroup to the other tree species) are recovered from these data. The accepted species tree, uniting humans and chimps, is the most frequent gene tree, with the two minor trees occurring in very similar frequencies, consistent with the action of only ILS during the divergence of these species (42); however, different studies have reported different frequencies for the three possible gene trees. For example, an early study that analysed 11945 gene trees (42) and a more recent study that analysed 1730 gene trees (41) found that 77% and 62% of gene trees respectively grouped humans and chimps, 12% and 20% respectively grouped chimps and gorillas, and 11% and 18% respectively grouped humans and gorillas. The discrepancies in these numbers reflect both the different data types and data quality available to each study, as well as differences in the methods used to reconstruct gene trees. However, both studies made the single-tree assumption for each individual gene locus; recombination within each locus violates this assumption. The MAST model avoids this assumption by using mixtures of trees—in principle, we expect that estimates of tree weights from the MAST model to be more accurate than estimates of gene tree frequencies from previous studies. Nonetheless, we still expect the MAST model to recover the highest weight for the tree grouping humans and chimps, and lower but approximately equal weights for the two minor trees.

**Fig. 4.**
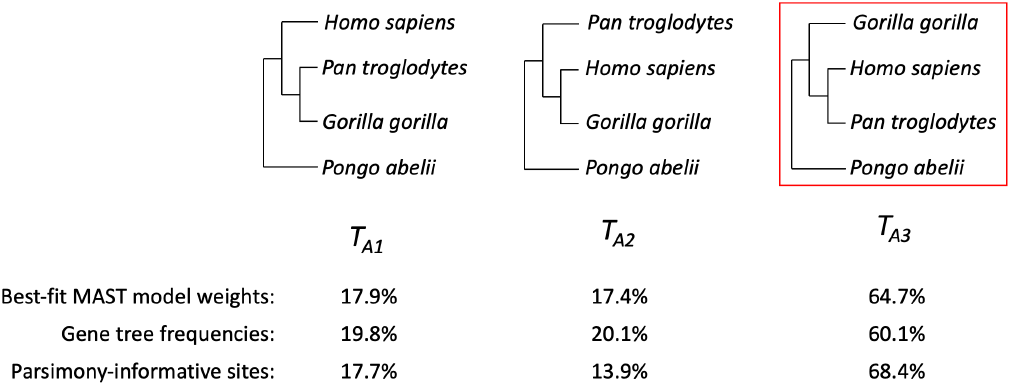
The three topologies for empirical dataset A. *T*_*A*3_ is the commonly accepted species tree.

The second empirical dataset (‘B’) includes three species from the genus *Macaca* (*M. fascicularis, M. mulatta, M. nemestrina*) and the mandrill (*Colobus angolensis palliatus*), a clade in which a previous analysis found substantial evidence for introgression between *M. nemestrina* and *M. fascicularis* (41). Thus, for this dataset we expect the MAST model to recover the highest weight for the accepted species tree uniting *M. fascicularis* and *M. mulatta* (*T*_*B*3_ in Figure 5), the second highest weight for the minor tree affected most by introgression (uniting *M. nemestrina* and *M. fascicularis*), and the lowest weight for the minor tree uniting *M. mulatta* and *M. nemestrina*.

**Fig. 5.**
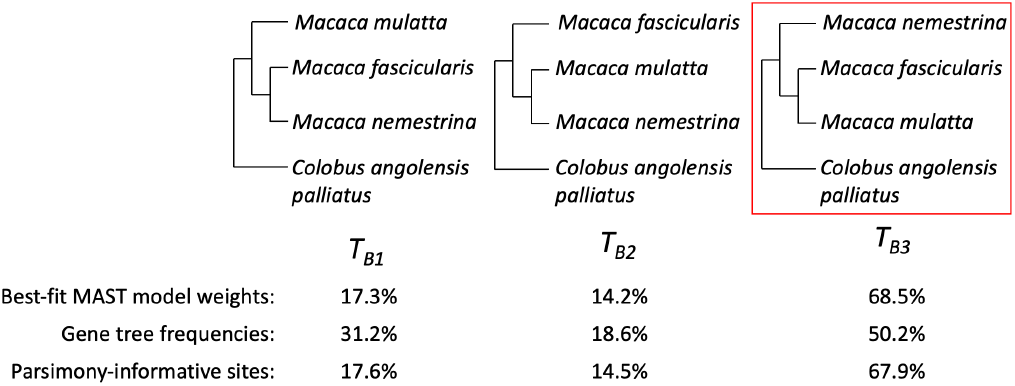
The three topologies for empirical dataset B. *T*_*B*3_ is the commonly accepted species tree.

The third empirical dataset (‘C’) contains the six species (human, chimp, gorilla, and the three *Macaca* species) that represent the ingroups from the first two datasets. Since we have a *priori* information which suggests that all three possible rooted trees are possible for each of these ingroups, we applied a MAST model with 9 trees (Supplementary Figure S5), where all three resolutions of each ingroup clade are paired with all three resolutions of the other ingroup clade. In principle, one should be able to draw similar conclusions from these 6-taxon datasets as one could from the two independent analyses of the four-taxon datasets by summing the relevant tree weights (see below).

The fourth empirical dataset (‘D’) focuses on the relationships among four Platyrrhine (“New World Monkey”) species: *Callithrix jacchus, Aotus nancymaae, Saimiri boliviensis*, and *Cebus capucinus imitator*, including *Maccaca mulatta* as an outgroup. There is disagreement about the species tree among the four focal taxa. Gene-tree-based analyses (41) support a caterpillar tree in which *Aotus* is the sister group to a clade uniting *Saimiri* and *Cebus* (*T*_*D*1_ in Supplementary Figure S6). However, concatenated ML analysis fails to recover this species tree, instead returning a symmetrical tree likely caused by a known inconsistency in ML methods when the underlying gene trees are highly discordant (43–45). The MAST model should in principle avoid statistical inconsistencies associated with the single-tree assumption because it explicitly accounts for the existence of multiple histories in an alignment. Thus, we sought to test the performance of the MAST model in this well-studied empirical test case. To do this, we applied a MAST model that included the three ingroup topologies that were most commonly found from the gene trees in a previous study (Supplementary Figure S6; (41)).

We analysed each empirical dataset using the same approach. First, we filtered the original 1730 locus dataset to retain only those loci that were present in all of the selected species, which resulted in each dataset containing approximately 1600 loci and around 1.6 million base pairs (Table 1). We analysed each dataset using standard single-tree concatenated ML analyses (using default settings in IQ-TREE2), as well as the six multitree mixture models described by the six submodels of the MAST model in Figure 2, using the trees topologies described above as the input topologies for the MAST model. Finally, to facilitate comparisons with other quantities of interest, we calculated for each of the input topologies: (1) the number of single-locus trees that matched each MAST tree, where each single locus tree was estimated with default parameters in IQ-TREE2; and (2) the total number of base pairs, (3) total number of variable sites, and (4) total number of parsimony informative sites in all of the single-locus alignments whose tree matched each MAST tree.

## Results

### Simulations 1-3: The MAST model performs well when the model is correctly specified, with or without introgression

Our extensive simulations demonstrate that the unlinked (Figure 6 and Supplementary Figure S2) and linked (Supplementary Figures S1 and S3) MAST models perform well when the model used for analysis matches that used to simulate the data. In both the linked and unlinked models, the error associated with all linked and unlinked models increases as the number of trees in the mixture increases, and as the number of tips in the tree decreases. This is expected, because in our simulations we held the total length of the alignment and the distribution of branch lengths constant. Thus, the amount of information available to estimate each parameter decreases (and thus the expected error increases) as the number of trees increases, and as the number of tips in each tree decreases. The key parameters of interest for the MAST models are the tree weights (top panel, Figure 6 and Supplementary Figure S1). In the best-case scenario (comprised of 2 trees, each of which contains 20 taxa) the RMSE of the tree weights was very low, at around 0.01 for both the unlinked and linked models. In the worst-case scenario (comprised of 10 trees, each of which contains 6 taxa) the error was much higher, at around 0.05 for both the unlinked and linked models, although this is still acceptably low in absolute terms. For both the unlinked and linked models the fit to data simulated under the matching MAST model is much better than for the mis-specified single-tree model (bottom panel, Figure 6 and Supplementary Figure S1); the improvement in the fit of the true model increases (i.e. the difference in BIC becomes more negative) as the number of trees and the number of tips in each tree increases. This is expected because a single-tree model becomes an increasingly poor fit to data simulated under more trees.

**Fig. 6.**
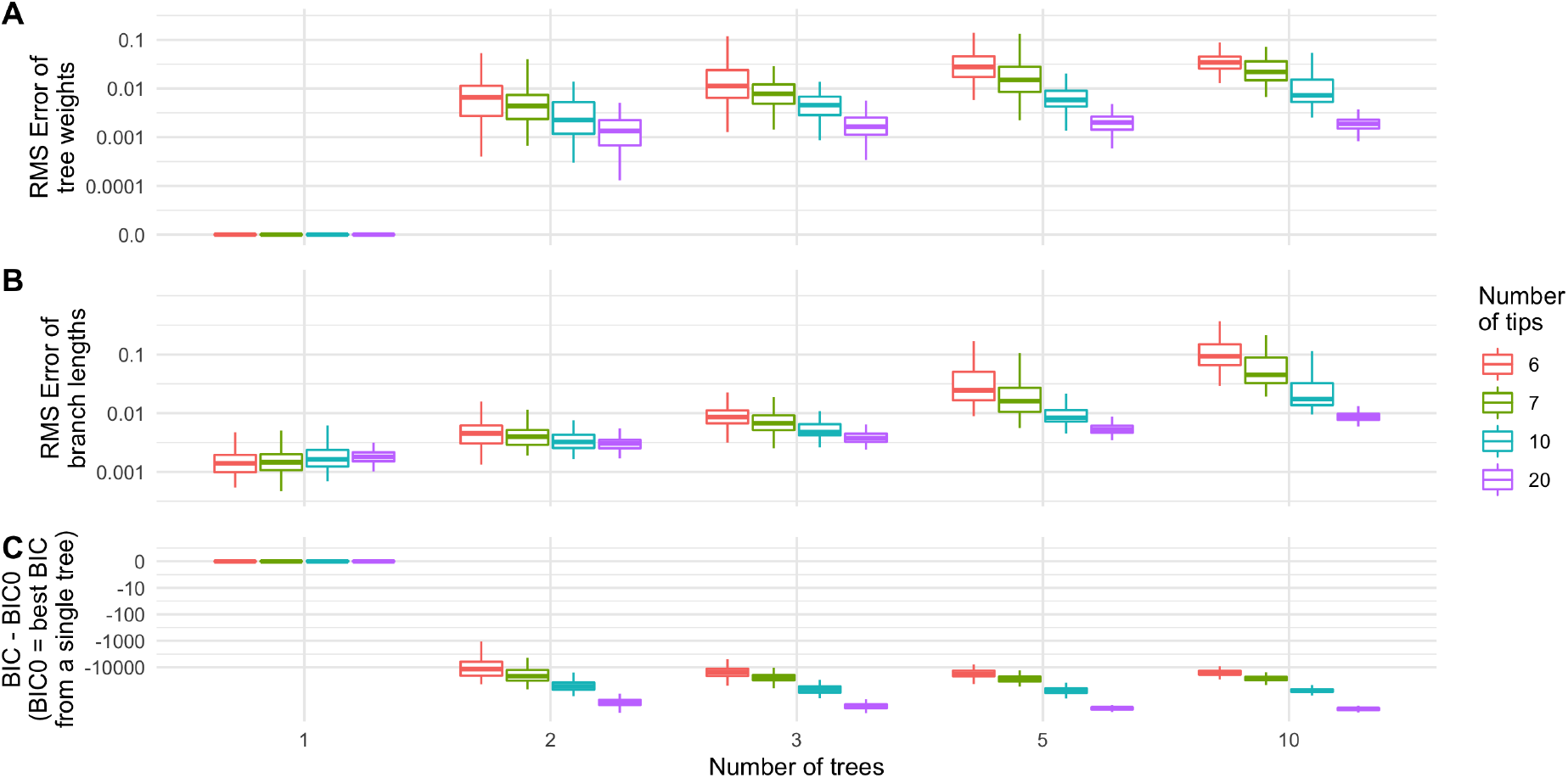
Performance of MAST model with unlinked parameters over all the trees (i.e. MAST submodel 1) on simulated data when the true topologies are given. All the trees have unlinked substitution matrices, nucleotide frequencies, and gamma parameters. The distribution of (A) the Root-Mean-Squared (RMS) error of the tree weights; (B) the RMS error of the branch lengths; and (C) *BIC* – *BIC*0, the difference between the BIC from the unconstrained MAST model (*BIC*) and that from a single-tree model (*BIC*0), for different numbers of trees with various numbers of tips. The negative values of BIC - BIC0 indicated that the MAST model is a better model compared with the standard single-tree model.

We also simulated scenarios with introgression, such that the minor trees are not expected to be equal in frequency. In these simulations *T*_*E*1_ is the species tree (Supplementary Figure S4) and increasing introgression makes topology *T*_*E*2_ increasingly frequent. When the introgression rate was between 0 and 0.6, *T*_*E*1_ is the optimal tree in the single-tree model (Figure 7A) and the tree with the highest weight in the MAST model (Figure 7B). When the introgression rate is above 0.6, in most datasets the single-tree model and the MAST model reported *T*_*E*2_ as the optimal tree and the topology with the highest tree weight, respectively. The results are as expected from the simulations that were carried out (i.e. the topology matching the introgressed history does in fact become the most common). The MAST model is a much better fit when the tree topologies *T*_*E*1_, *T*_*E*2_ are more equal in frequency, though it is a better fit across all of parameter space (because there is always ILS; Figure 7C).

**Fig. 7.**
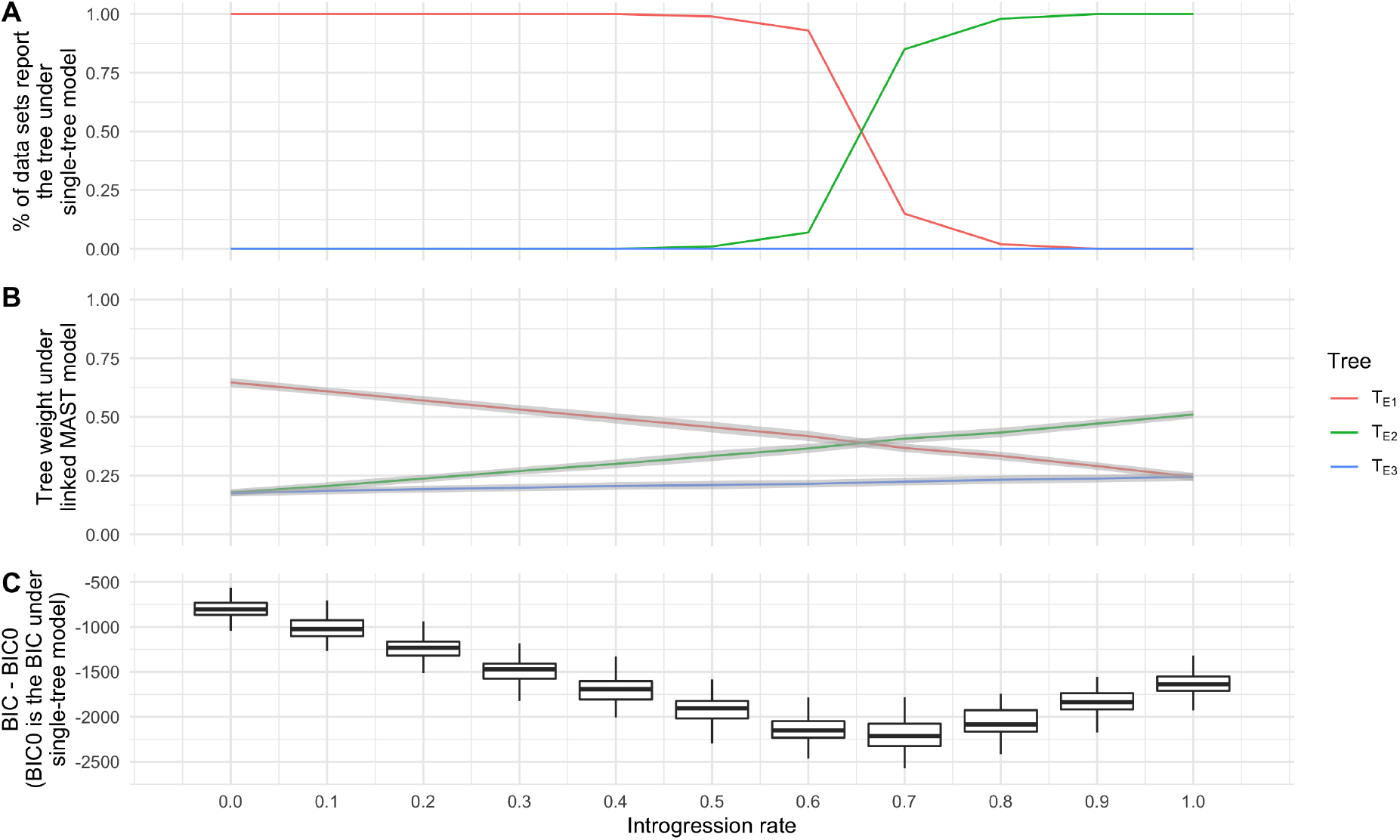
Results from simulated datasets with introgression rate *r* ∈ {0.0,0.1,…, 1.0} (A) Fitting the concatenated alignment under a single-tree model. The most likely tree topology becomes *T*_*E*2_ at very high rates of introgression; (B) The tree weights under the linked MAST model. The coloured lines indicate the mean and the grey regions indicate the standard deviation among the 100 datasets; (C) *BIC* – *BIC*0, the difference between the BIC from the linked MAST model (*BIC*) and from a single-tree model (*BIC*0).

### Simulation 4: The MAST model is robust to the inclusion of trees with no support in the underlying data

To test the robustness of the MAST model to the inclusion of incorrect additional topologies, we simulated data under a single topology but fit the data under a MAST model with 10 topologies. Including up to nine trees that have no support in the data had surprisingly little effect on the conclusions. Figure 8A reveals that the true tree (which was always one of the trees included in the MAST model) had the maximum weight among all of the trees included in the MAST model in the majority of simulations, regardless of the simulation conditions. Importantly, the inclusion of incorrect trees in the MAST model always led to large increases in the BIC score, such that researchers using this method to select the best model would reject the additional trees, and instead prefer the results from a single-tree model (Figure 8B).

**Fig. 8.**
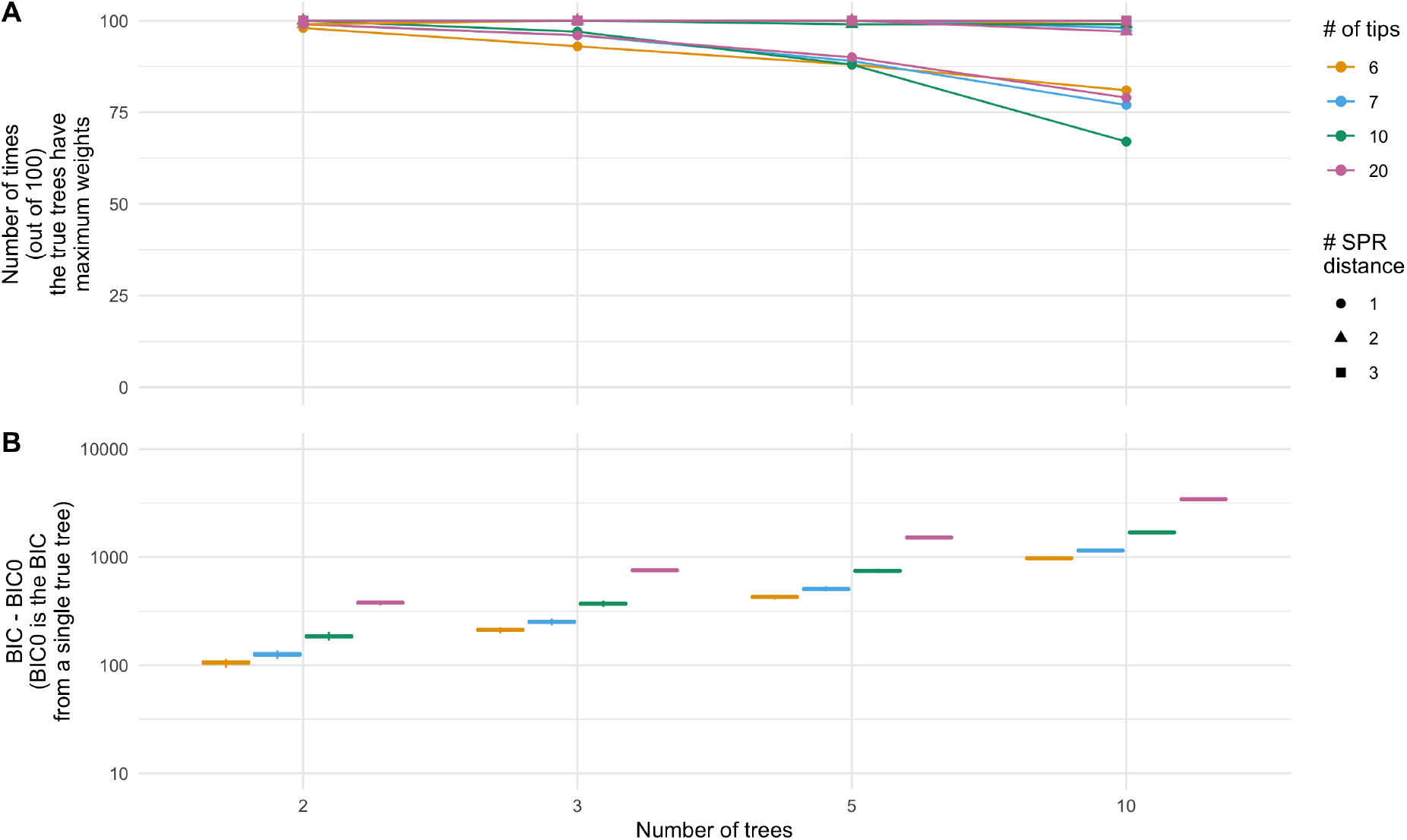
Performance of MAST model on simulated data when it was applied to the datasets simulated under a single tree. We used an unconstrained MAST model in which each class has its own substitution matrices, base frequencies, and rate heterogeneity model. (A) The percentage of times the true trees have the maximum weights for different numbers of input tree topologies and various SPR distances between the actual topology and the other input topologies. (B) *BIC* – *BIC*0, the difference between the BIC from the MAST model (*BIC*) and those from a single-tree model (*BIC*0).

These simulations also reveal some of the fundamental limitations of the MAST model to distinguish among very similar trees. When incorrect trees included in the MAST model were sufficiently different from the true tree (i.e. when the SPR distance of each incorrect tree in the mast model was 2 or 3 SPR moves from the true tree), the percentage of simulations for which the true tree had the maximum weight remained near 100% regardless of the other simulation conditions. However, when the incorrect trees included in the MAST model were close to the true tree (i.e. when they differed from the true tree by a single SPR move), the percentage of simulations for which the true tree had the maximum weight dropped to around 70% in the worst case (trees with 10 tips; Figure 8A). This general trend is expected, because more similar trees will share more branches in common, making it more difficult for any model to distinguish between them. Our results therefore help to quantify some of the analytical limits of multitree mixture models as currently implemented.

### Empirical dataset A: Incomplete lineage sorting in the Great Apes

Figure 4 shows the three possible tree topologies *T*_*A*1_, *T*_*A*2_, *T*_*A*3_ for empirical dataset A, which is made up of four Great Apes (Table 1). We applied multiple methods to these alignments in order to estimate the frequency of the three tree topologies. Single-tree analyses applied to each gene separately reported 19.8%, 20.1%, and 60.1% of the genes with topologies *T*_*A*1_, *T*_*A*2_, *T*_*A*3_, respectively (Figure 4; Supplementary Table S1). All MAST submodels reported similar tree weights of 17.9%, 17.4%, and 64.7% (Figure 4; Table 2). All methods find that the topology uniting human and chimpanzee is the most common, with the two minor topologies having approximately equal weights; these results are as expected from all previous analyses.

**Table 2.**
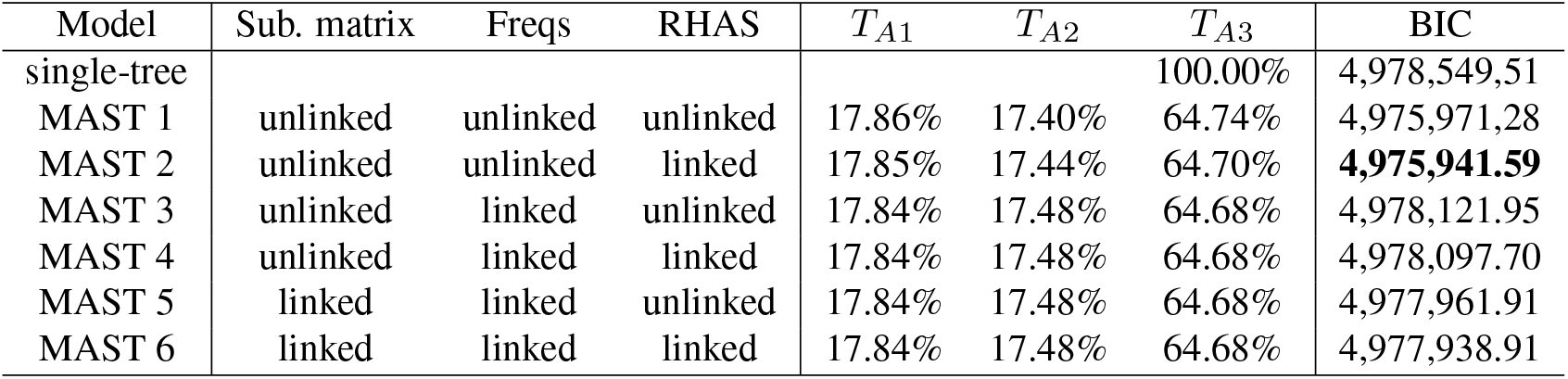
Results of the empirical dataset A when applying IQ-TREE with a standard single-tree model and different MAST submodels with GTR+G substitution model. There are six submodels of MAST, representing different combinations of linked or unlinked substitution matrix (2nd column), nucleotide frequencies (3rd column), and rate heterogeneity across sites (4th column). The 5th-7th columns are the weights of the trees *T*_*A*1_, *T*_*A*2_, *T*_*A*3_. The 8th column lists the BIC values of different models. The bolded figure is the best BIC value which is from the MAST submodel 2.

The proportions of different topologies estimated by MAST are closer to the proportions estimated for individual nucleotide sites than for the corresponding gene trees than the percentage of the gene trees (Supplementary Table S1). This may be because the weights of the MAST model more closely approximate the percentage of the sites (instead of the percentage of genes) belonging to different topologies. The BIC score from MAST submodel 2 was the best (Table 2), indicating that the MAST model with unlinked substitution model, unlinked frequencies and linked RHAS was the best model among different MAST submodels for this dataset.

Regardless, the BIC values of all MAST submodels were much lower than the BIC value reported by the single-tree model (Table 2), showing that a multitree-mixture model had a much better fit to the data.

### Empirical dataset B: Introgression in *macaques*

Figure 5 shows the three possible tree topologies *T*_*B*1_, *T*_*B*2_, *T*_*B*3_ for empirical dataset B, which is made up of multiple macaque species. Analyses of the individual gene trees using single-tree models for each revealed a large asymmetry in minor topologies (31.2%, 18.6%, and 50.2% for *T*_*B*1_, *T*_*B*2_, *T*_*B*3_ respectively; Supplementary Table S2). However, both the proportions of parsimony-informative sites (17.6%, 14.5%, and 67.9% for *T*_*B*1_, *T*_*B*2_, *T*_*B*3_ respectively; Supplementary Table S2) and the weights from the different MAST submodels (all around 17.3%, 14.2%, 68.6% for *T*_*B*1_, *T*_*B*2_, *T*_*B*3_ respectively; Figure 5; Table 3) showed much more similar proportions and weights for the minor trees. Although the minor trees are still substantially different in frequency using the MAST analysis—consistent with introgression in this clade—the difference between them is much lower. As in empirical dataset A, this result indicates that inferences directly from gene trees may be underestimates of the frequency of the most common tree, and overestimates of the frequencies of the minor trees.

**Table 3.** Results of the empirical dataset B when applying IQ-TREE with a standard single-tree model and different MAST submodels with GTR+G substitution model. There are six submodels of MAST, representing different combinations of linked or unlinked substitution matrix (2nd column), nucleotide frequencies (3rd column), and rate heterogeneity across sites (4th column). The 5th-7th columns are the weights of the trees *T*_*B*1_, *T*_*B*2_, *T*_*B*3_. The 8th column lists the BIC values of different models. The bolded figure is the best BIC value, which is MAST submodel 2.

| Model | Sub. matrix | Freqs | RHAS | $T_{B1}$ | $T_{B2}$ | $T_{B3}$ | BIC |
| --- | --- | --- | --- | --- | --- | --- | --- |
| single-tree |  |  |  |  |  | 100.00% | 4,906,941.36 |
| MAST 1 | unlinked | unlinked | unlinked | 17.29% | 14.15% | 68.55% | 4,905,832.06 |
| MAST 2 | unlinked | unlinked | linked | 17.29% | 14.19% | 68.52% | **4,905,808.79** |
| MAST 3 | unlinked | linked | unlinked | 17.27% | 14.24% | 68.49% | 4,906,632.17 |
| MAST 4 | unlinked | linked | linked | 17.27% | 14.25% | 68.48% | 4,906,605.01 |
| MAST 5 | linked | linked | unlinked | 17.27% | 14.24% | 68.50% | 4,906,651.67 |
| MAST 6 | linked | linked | linked | 17.27% | 14.23% | 68.50% | 4,906,633.71 |

### Empirical dataset C: Great Apes + *macaques*

Supplementary Figure S5 shows nine tree topologies for the empirical dataset C. This dataset combines the ingroup taxa from empirical datasets A and B, allowing us to test the accuracy of MAST when there are more possible topologies: the nine topologies represent every combination of the three topologies present in each of empirical datasets A and B. The frequencies of the nine tree topologies were similar across gene trees and sites in standard analysis (Supplementary Table S3) as well as largely similar to the results across MAST submodels (Table 4). MAST submodels 1 and 2 are the two best-fit models to the dataset according to the BIC values (Table 4), and both give tree weights that are quite close to the corresponding tree weights for the respective analyses in empirical datasets A and B (Supplementary Tables S4 and S5).

**Table 4.** Results of the empirical dataset C when applying IQ-TREE with a standard single-tree model and different MAST submodels with GTR+G substitution model. Six submodels of MAST are for different combinations of linked or unlinked substitution matrix, nucleotide frequencies, and rate heterogeneity across sites. The 2nd - 10th columns are the estimated tree weights between the topologies *T*_*C*1_, *T*_*C*2_,…, and *T*_*C*9_ for different MAST submodels. The bolded figure is the best BIC value among different submodels.

| Model | $T_{C1}$ | $T_{C2}$ | $T_{C3}$ | $T_{C4}$ | $T_{C5}$ | $T_{C6}$ | $T_{C7}$ | $T_{C8}$ | $T_{C9}$ | BIC |
| --- | --- | --- | --- | --- | --- | --- | --- | --- | --- | --- |
| single-tree |  |  |  |  |  |  |  |  | 100.0% | 5,187,194.8 |
| MAST 1 | 0.4% | 7% | 8.4% | 7.7% | 2.9% | 18.3% | 13% | 8.7% | 33.6% | **5,183,982.5** |
| MAST 2 | 0.4% | 10.4% | 8.2% | 2.1% | 2.5% | 14% | 13.1% | 8.4% | 41.1% | 5,183,988.4 |
| MAST 3 | 0.2% | 8% | 5.2% | 1.1% | 0.2% | 17.4% | 15.2% | 2.4% | 50.4% | 5,186,041.4 |
| MAST 4 | 0.2% | 0.2% | 3.9% | 0.6% | 0.8% | 29.3% | 12.7% | 19.8% | 32.5% | 5,185,924.7 |
| MAST 5 | 0.0% | 0.8% | 9.8% | 1.9% | 0.4% | 18.2% | 17.1% | 11.3% | 40.4% | 5,186,243.3 |
| MAST 6 | 0.0% | 0.7% | 11.1% | 1.9% | 1.8% | 20.7% | 19.3% | 8.4% | 36% | 5,186,194.1 |

### Empirical dataset D: Overcoming known biases in concatenated maximum likelihood

As mentioned, maximum likelihood has a known bias toward symmetrical trees (i.e. *T*_*D*3_ in Supplementary Figure S6) when there is a large amount of underlying discordance and the true species tree is asymmetrical (i.e. *T*_*D*1_ or *T*_*D*2_ in Supplementary Figure S6). Indeed, when analyzed under ML using a single-tree model, data from four Platyrrhine monkeys support a symmetrical tree (Table 5). In contrast, counts of genes trees and parsimony-informative sites support the asymmetrical tree *T*_*D*1_ as the species tree (Supplementary Table S6). Similarly, analyses using the MAST submodels also tended to return *T*_*D*1_ as the topology with the highest weight (Table 5). Among all the models, the MAST submodel 2 had the best BIC value, with reported tree weights 42.4%, 28.1%, 29.6% for the topologies *T*_*D*1_, *T*_*D*2_, *T*_*D*3_. The tree weights are similar to the proportions of parsimony-informative sites supporting each of these topologies (i.e. 36.7%, 32.2%, 31.1%; Supplementary Table S6). It is notable that two MAST models estimated different trees with the highest weights (submodels 3 and 4; Table 5), though submodel 2 has a much lower BIC value than either of these. Overall, these results suggest that the MAST model is able to analyse a concatenated alignment using maximum likelihood, but without the biases that come with the single-tree assumption.

**Table 5.** Results of the empirical data D when applying IQ-TREE with a standard single-tree model and different MAST submodels with GTR+G substitution model. Six submodels of MAST are for different combinations of linked or unlinked substitution matrix (2nd column), nucleotide frequencies (3rd column), and rate heterogeneity across sites (4th column). The 5th, 6th, and 7th columns are the estimated tree weights between the topologies *T*_*D*1_, *T*_*D*2_, and *T*_*D*3_ for different MAST submodels, respectively. The bolded figure is the best BIC value among different submodels.

| Model | Sub. matrix | Freqs | RHAS | $T_{D1}$ | $T_{D2}$ | $T_{D3}$ | BIC |
| --- | --- | --- | --- | --- | --- | --- | --- |
| single-tree |  |  |  |  |  | 100.00% | 6,185,094.0 |
| MAST 1 | unlinked | unlinked | unlinked | 40.3% | 23.0% | 36.8% | 6,177,609.0 |
| MAST 2 | unlinked | unlinked | linked | 42.4% | 28.1% | 29.6% | **6,177,535.7** |
| MAST 3 | unlinked | linked | unlinked | 3.5% | 4.7% | 91.8% | 6,182,942.1 |
| MAST 4 | unlinked | linked | linked | 2.1% | 81.3% | 16.7% | 6,182,954.3 |
| MAST 5 | linked | linked | unlinked | 42.4% | 32.0% | 25.6% | 6,184,689.7 |
| MAST 6 | linked | linked | linked | 42.4% | 32.0% | 25.5% | 6,184,618.7 |

## Discussion

We have introduced the mixtures across sites and trees (MAST) model, which assumes that every site in a concatenated alignment may have evolved from a mixture of trees. This flexible assumption allows the method to be applied to the alignments that include multiple tree topologies, which is presumably true of almost any large dataset from a recombining genome. The implementation of the method allows different combinations of linked and unlinked parameters when estimating the substitution matrix, nucleotide or amino acid frequencies, and the rate heterogeneity across sites (RHAS) across different trees. This flexibility allows researchers to have many of the advantages of concatenated analyses—e.g. a large amount of data and accurate estimate of complex substitution processes—while still incorporating gene tree heterogeneity, but without the need to make assumptions about the existence and location of putatively non-recombining loci. As such, the MAST model opens up the opportunity to study topological discordance in deep time, past the point where information from small, non-recombining gene tree alignments can be informative about relationships (31).

From the simulation experiments we carried out it appears that the estimates of the parameters are relatively reliable, especially when the number of taxa is large or the alignments are long, relative to the number of trees included in the MAST model. Although only the MAST model with unlinked parameters (i.e. submodel 1) and the MAST model with linked parameters (i.e. submodel 6) were tested, these two models are the models with the most and the least number of parameters, respectively, and therefore should encompass the accuracy of models between them. The overall results indicate that the parameters are identifiable. The identifiability of parameters in complex models, like mixture models, has been addressed previously (26, 46). Some research has shown strong theoretical evidence that, when mixture models are applied, cases where trees are non-identifiable are rare (46–48). Rhodes and Sullivant (2012) gave an upper bound on the number of classes that ensures the generic identifiability of trees in models with a multi-tree mixture. Their method was based on the mixtures from different trees, provided that all the topologies share a certain type of common substructure in which a tripartition *A*|*B*|*C* exists such that the splits *A*|*B* ∪ *C* and *A* ∪ *C*|*B* are compatible with all trees. Parameters in the multi-tree mixture model are generically identifiable provided *m* < *k*^*j*–1^ where *m* is the number of classes, *k* is the number of states (i.e. 4 for nucleotides; 20 for amino acids), and the number of taxa in the partition *A* and in the partition *B* are both greater than or equal to *j*. However, establishing the identifiability of model parameters when there is no commonality between the trees remains an open problem (46).

In order to use the MAST model to perform an analysis, the user must input a set of pre-specified tree topologies. A rooted three-taxon tree has only three possible topologies, but the number of topologies goes up super-exponentially with the number of tips (Table 3.1 in (49)). This fact means that it will usually not be feasible to specify all possible topologies that exist in a moderate-sized dataset; for example, in empirical dataset D we only studied 3 of 15 topologies. While this would seem to limit the applicability of MAST, often there will be a smaller set of topologies that are relevant to any particular study. For instance, even in a tree with 100 species, it may be the relationships among a smaller number of clades that are relevant: if ILS only occurs on one branch of the tree, then there are still only three relevant alternative topologies, no matter the number of tips. In general, we recommend that users specify known alternative hypotheses—or carry out an exploratory analysis of individual gene trees—in order to choose a manageable set of topologies as input to the MAST model.

There are multiple known biases when carrying out concatenated analyses under the “treelikeness” assumption. As mentioned in the Introduction, single-tree concatenated maximum likelihood is statistically inconsistent in the presence of large amounts of discordance: it will return the incorrect tree with increasing probability as more data are added (43). Our analyses of Platyrrhine monkeys suggest that the MAST model can solve this problem, giving the highest weight to the topology favored by other (statistically consistent) methods. In addition to inferring the wrong tree topology, the branch lengths inferred from concatenated analyses are biased in the presence of discordance (21, 50). Such biases can lead to misestimation of divergence times when using the entire concatenated alignment. The MAST model allows researchers to estimate the branch lengths of individual topologies—we therefore recommend estimating divergence times using branch lengths obtained from the topology matching the species tree. While these times still represent genic divergence (and not species divergence; (51)), they will be free of the bias associated with single-tree concatenation.

The output of our method is a set of weights associated with each input tree topology. Although the MAST model is not based on a particular biological model of discordance (e.g. the MSC or MSNC), we expect that the estimated weights should correspond to biologically relevant features of the data. Both our analyses of simulated and empirical data revealed that the reported weights in the MAST model are highly correlated with the proportion of phylogenetically informative sites which support each tree. This correlation is expected because the likelihood of each site is calculated as the weighted sum of the likelihood of the site over all the trees and the overall likelihood value is the product of the likelihoods over all the sites. This result, together with the accurate estimation of minor tree weights, means that we can use these estimates to infer introgression from MAST output. Common tests for introgression are based on the expectation that the two minor trees are equal in frequency (e.g. the “ABBA-BABA” test; (52)). One post hoc approach to inferences of introgression using MAST would be to test for a significant difference in the weights supporting each of two minority trees. Alternatively, it should be possible to compare the likelihoods of models that either link or unlink the weights of the minority trees. Greater support for the unlinked model would indicate that the two trees are not equal in frequency, and would support an inference of introgression. Such an approach would be of great benefit to testing for introgression deeper in time, where individual phylogenetically informative sites and individual gene trees may not be accurate enough to make strongly supported inferences about introgression (41).

Finally, the MAST model can be used to assign posterior probabilities of membership in a class (i.e. topology) to each nucleotide site in an alignment. Because recombination will not completely erase spatial information about local topologies along a chromosome, these posterior probabilities should allow us to infer the locations of switches between topologies. A hidden Markov model (e.g. (53)) or similar approach can then be run to identify individually nonrecombining blocks of a longer alignment that contain only a single topology. Such an approach would also enable us to detect topology-switches in any type of alignment. For example, there is no barrier to applying MAST to an alignment of paralogous sequences, either to detect the presence of multiple different topologies or the location of changes in topology (i.e. ectopic gene conversion; (54)).

The MAST model is a flexible phylogenetic approach that allows each site in an alignment to have evolved from a mixture of trees. Each tree has its own topology, a separate set of branch lengths, a substitution model, a set of nucleotide or amino-acid frequencies, and a rate heterogeneity model. However, there are still some limitations to the current implementation. In addition to several future directions mentioned above, we would like to extend the MAST model to: (1) Perform a tree topology search for an input number of trees. This would relax the requirement that the user must pre-specify topologies; (2) Be able to compute the optimal number of trees needed to represent the input dataset. This would relax the requirement that the user specify the number of trees ahead of time; and (3) Find the best set of substitution models and RHAS models for each tree separately. This would allow much more flexibility in the range of evolutionary variation that can be accommodated. These directions are challenging but will be useful in analysing genome-scale datasets at any evolutionary timescale.

## Acknowledgements

We thank Fred Jaya, Huaiyan Ren, James Barbetti, Jeremias Ivan, Nhan Trong Ly, and Rahil Vora (Australian National University) for participating in our group meetings where these results were discussed and providing comments on the manuscript. This work was supported by an Australian Research Council Discovery Project (DP200103151); a U.S. National Science Foundation (DEB-1936187 to M.W.H); a Moore-Simons Foundation grant (https://doi.org/10.46714/735923LPI to B.Q.M.); and a Chan-Zuckerberg Initiative Grant for Essential Open Source Software for Science to B.Q.M. and R.L.

## Availability of software and supplementary materials

Data, scripts, and supplementary materials are available from the Dryad Digital Repository: http://dx.doi.org/10.5061/dryad.mgqnk992p.

MAST model has been implemented in IQ-TREE2, which is available in the Github: https://github.com/iqtree/iqtree2/releases/tag/v2.2.0.7.mx.

## Supplementary Figures and Tables

**Fig. S1.**
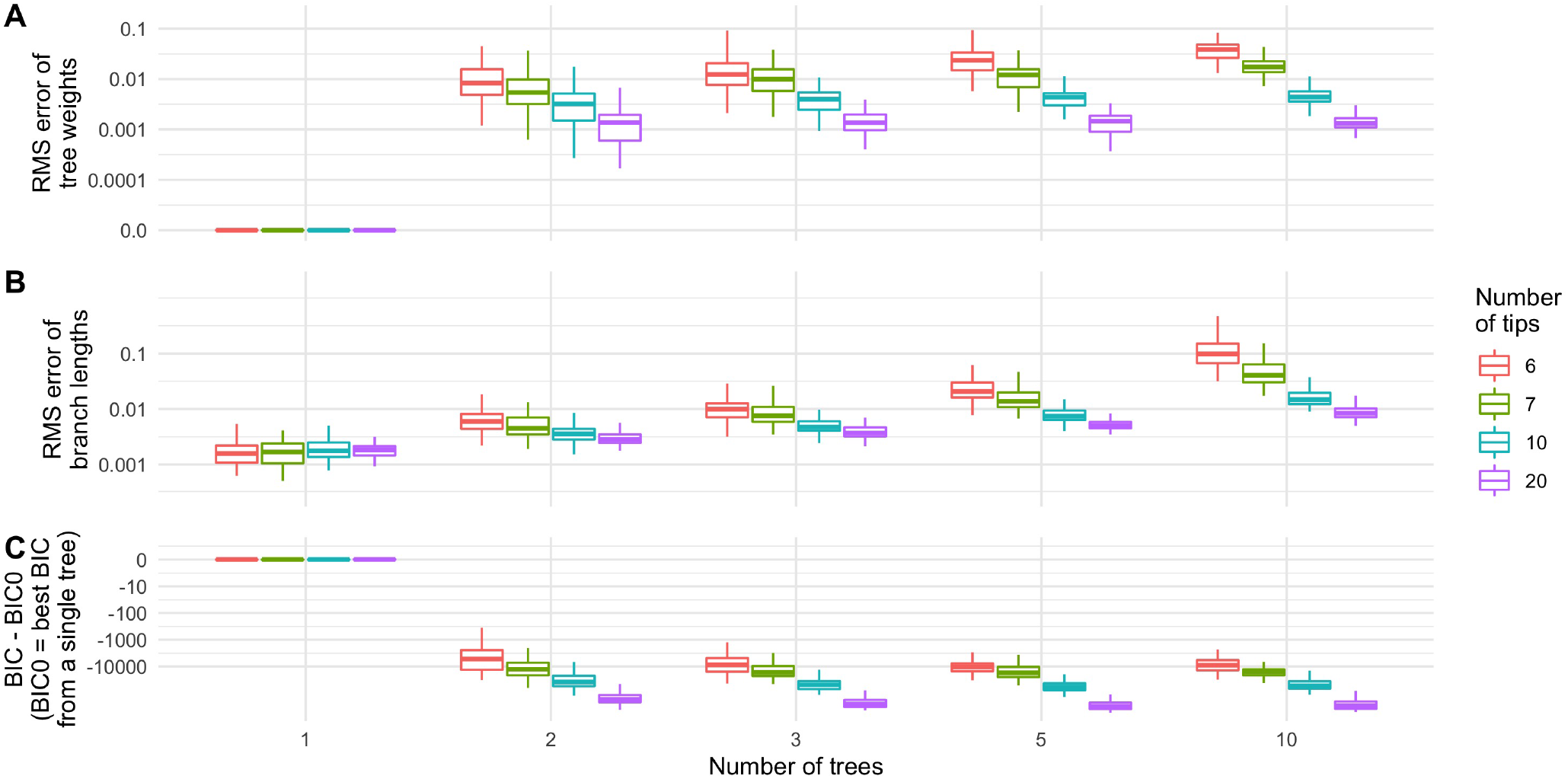
Performance of MAST model with linked parameters over all the trees (i.e. MAST submodel 6) on simulated data when the true topologies are given. All the trees share the same substitution matrices, nucleotide frequencies, and gamma parameters. The distribution of (A) the RMS error of the tree weights; (B) the RMS error of the branch lengths; and (C) *BIC* – *BIC*0, the difference between the BIC from the MAST model (*BIC*) and that from a single-tree model (*BIC*0), for different numbers of trees with various numbers of tips. The negative value of *BIC* – *BIC*0 indicated that the MAST model is a better model compared with the standard single-tree model.

**Fig. S2.**
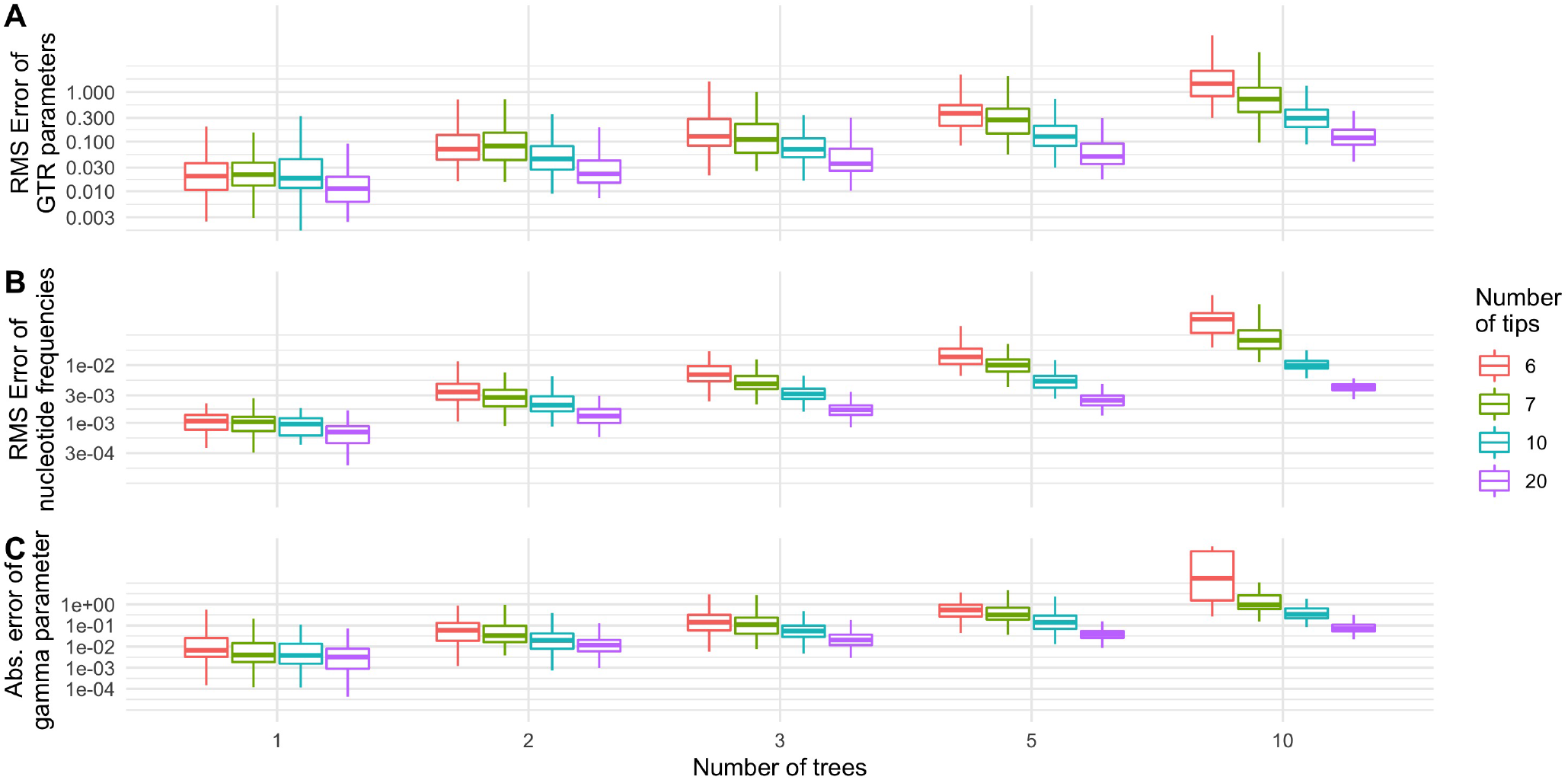
Performance of MAST model with unlinked parameters over all the trees on simulated data when the true topologies are given. All the trees have unlinked substitution matrices, nucleotide frequencies, and gamma parameters. The root-mean-squared (RMS) error of (A) the parameters of the GTR model, (B) the nucleotide frequencies, and (C) the gamma parameters, for different numbers of trees with various numbers of tips.

**Fig. S3.**
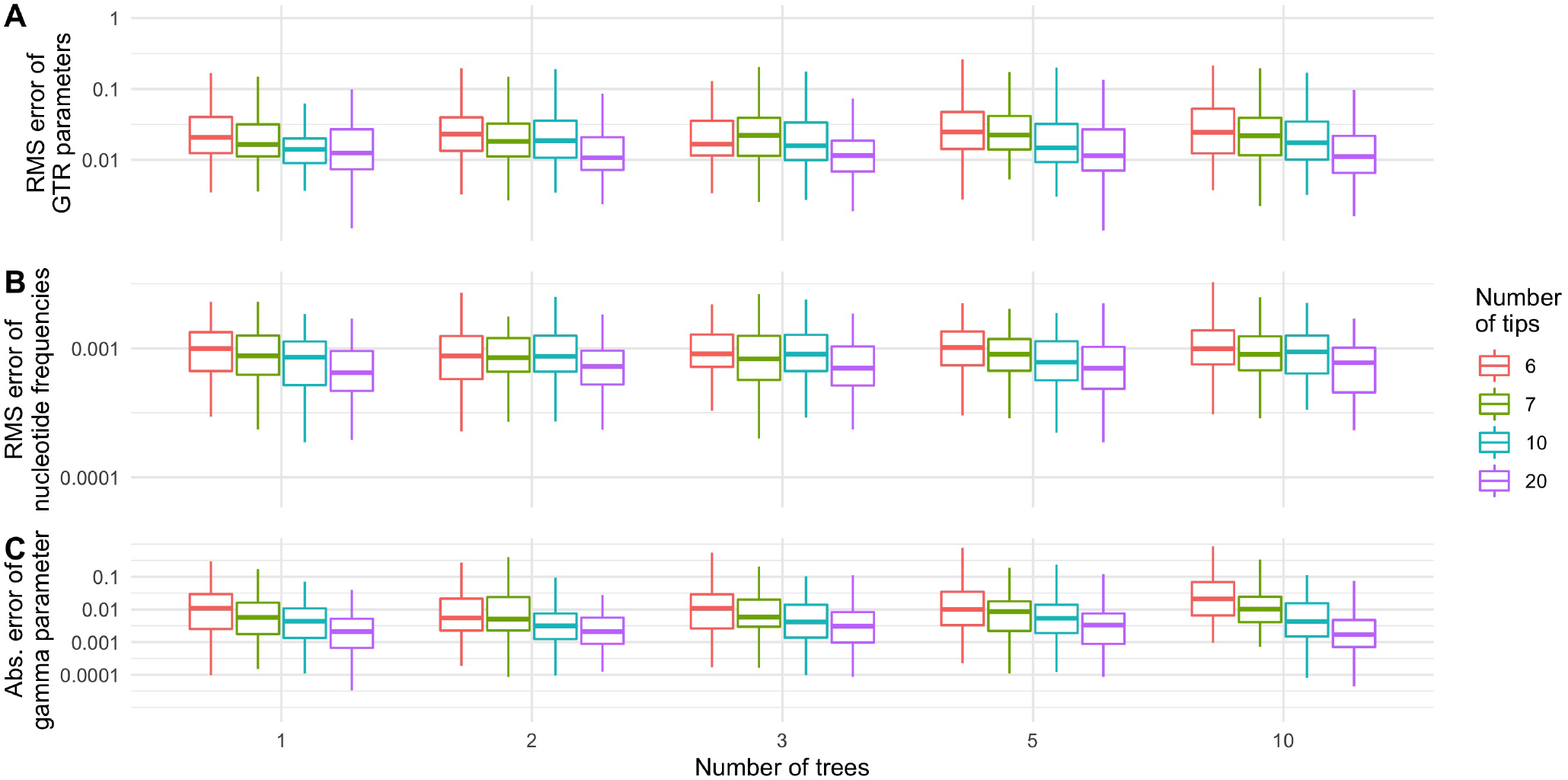
Performance of MAST model with linked parameters over all the trees on simulated data when the true topologies are given. All the trees share the same substitution matrices, nucleotide frequencies, and gamma parameters. The root-mean-squared (RMS) error of (A) the branch lengths, (B) the nucleotide frequencies, (C) the gamma parameters, for different numbers of trees with various numbers of tips.

**Fig. S4.**
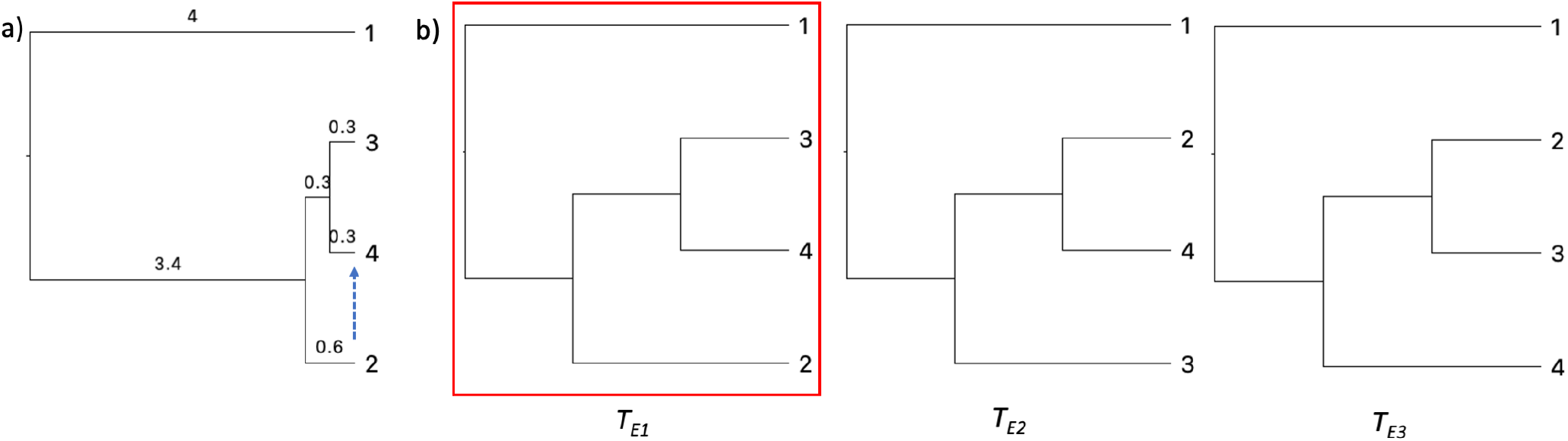
(A) Gene trees with introgression rate *r* ∈ {0.0,0.1,…, 1.0} were simulated in the direction from lineages 2 to 4 by *ms* under the coalescent model. The edge lengths are in coalescent time; (B) Three possible topologies for the simulated introgression dataset. *T*_*E*1_ is the species tree while some sites may evolve under the topologies *T*_*E*2_ and *T*_*E*3_ due to the introgression and incomplete lineage sorting.

**Table S1.** Detailed information of the genes in empirical dataset A. *T*_*A*3_ is the commonly accepted species tree.

| Topology | Number of genes<br>(Gene tree %) |  | Sum of gene lengths |  | Total number of<br>variable sites |  | Total number of<br>parsimony<br>informative sites |  |
| --- | --- | --- | --- | --- | --- | --- | --- | --- |
| $T_{A1}$ | 316 | 19.8% | 302,416 | 18.7% | 7,557 | 19.9% | 430 | 17.7% |
| $T_{A2}$ | 320 | 20.1% | 278,066 | 17.2% | 6,259 | 16.5% | 339 | 13.9% |
| $T_{A3}$ | 959 | 60.1% | 1,038,024 | 64.1% | 24,136 | 63.6% | 1,662 | 68.4% |
| Total | 1,595 |  | 1,618,506 |  | 37,952 |  | 2,431 |  |

**Fig. S5.**
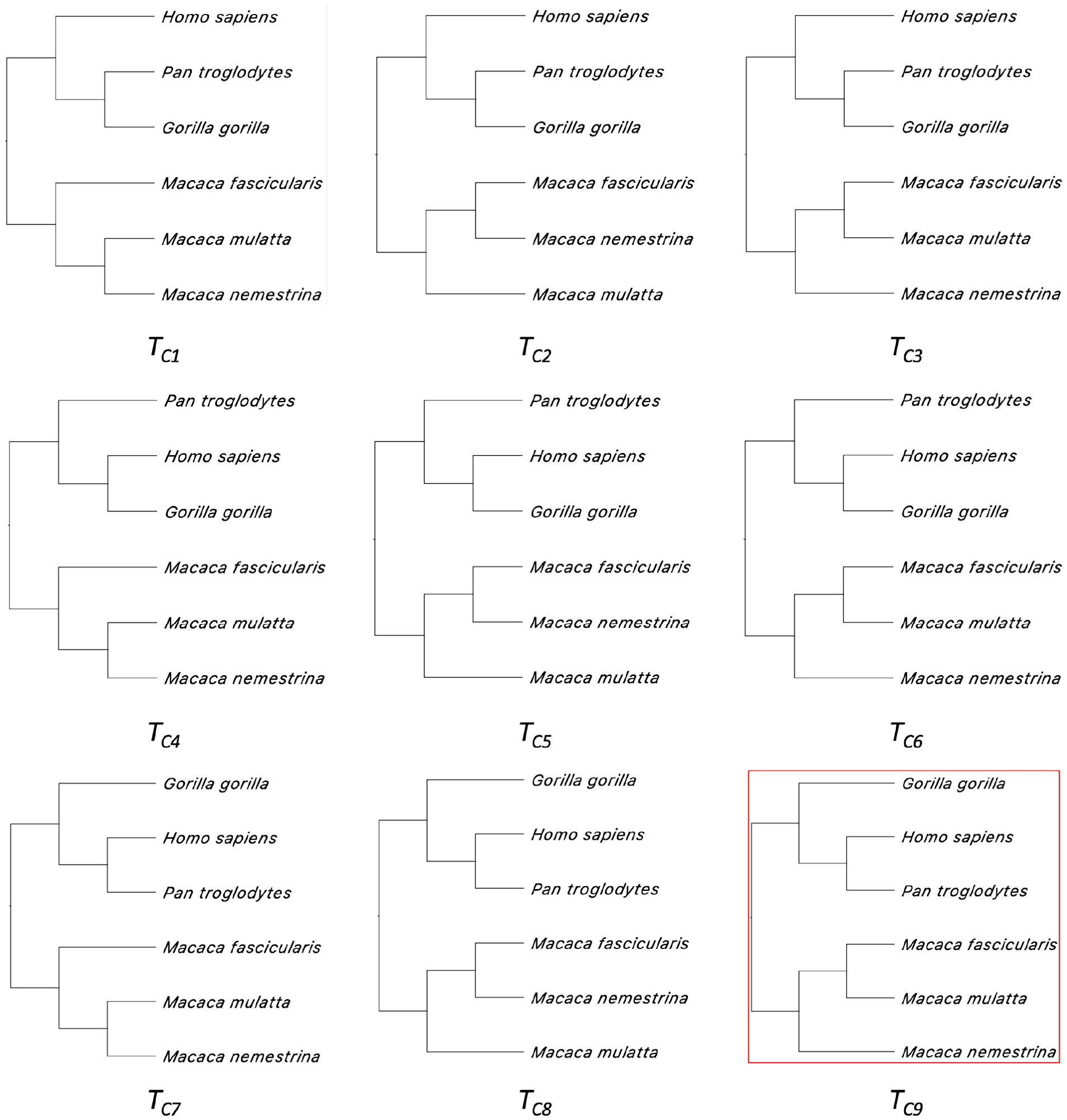
Nine topologies for empirical dataset C. *T*_*C*9_ is the commonly accepted species tree.

**Table S2.** Detailed information of the genes in empirical dataset B. *T*_*B*3_ is the commonly accepted species tree.

| Topology | Number of genes<br>(Gene tree %) |  | Sum of gene lengths |  | Total number of<br>variable sites |  | Total number of<br>parsimony<br>informative sites |  |
| --- | --- | --- | --- | --- | --- | --- | --- | --- |
| $T_{B1}$ | 499 | 31.2% | 438,469 | 26.9% | 8,218 | 26.6% | 214 | 17.6% |
| $T_{B2}$ | 298 | 18.6% | 298,016 | 18.3% | 5,817 | 18.8% | 176 | 14.5% |
| $T_{B3}$ | 802 | 50.2% | 892,678 | 54.8% | 16,857 | 54.6% | 824 | 67.9% |
| Total | 1,599 |  | 1,629,163 |  | 30,892 |  | 1,214 |  |

**Fig. S6.**
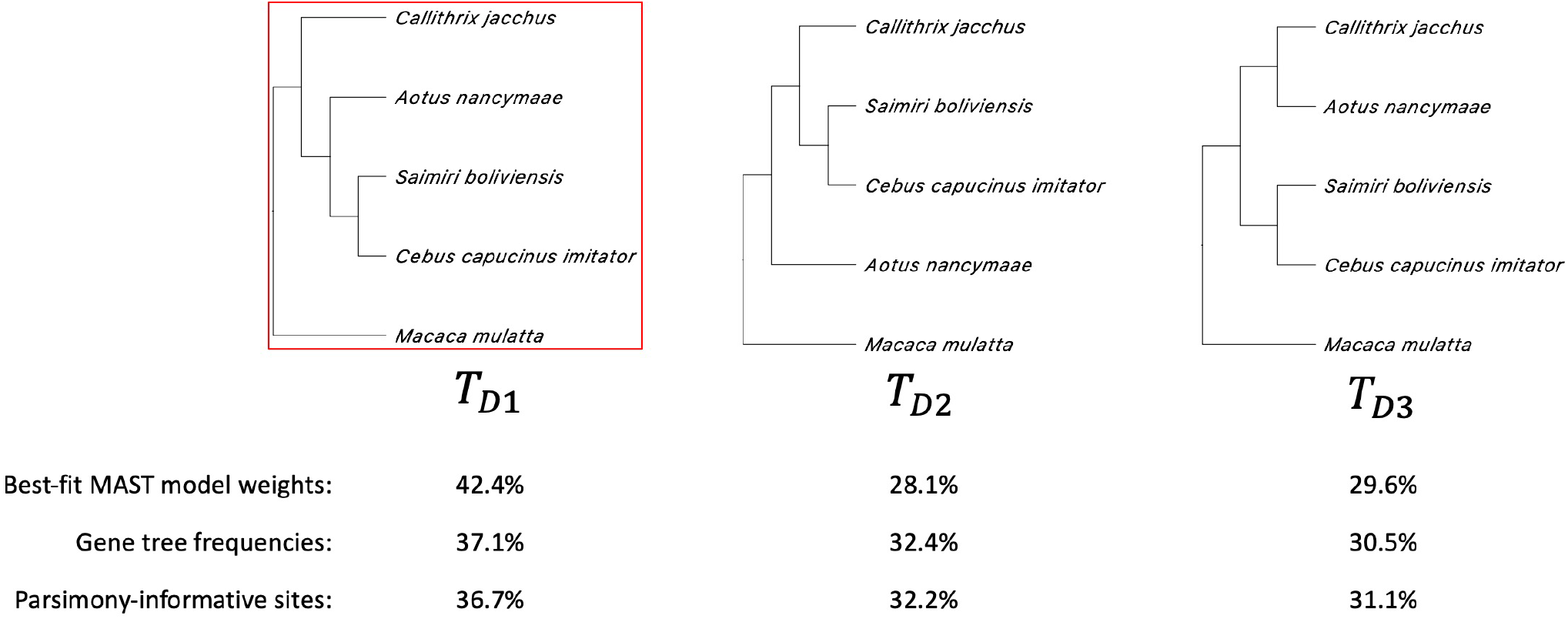
Three topologies for empirical dataset D. *T*_*D*1_ is the commonly accepted species tree.

**Table S3.** Detailed information of the genes in empirical dataset C. *T*_*C*9_ is the commonly accepted species tree.

| Topology | Number of genes<br>(Gene tree %) |  | Sum of gene lengths |  | Total number of<br>variable sites |  | Total number of<br>parsimony<br>informative sites |  |
| --- | --- | --- | --- | --- | --- | --- | --- | --- |
| $T_{C1}$ | 53 | 3.4% | 45,687 | 2.9% | 2,014 | 3.2% | 1,216 | 3.0% |
| $T_{C2}$ | 102 | 6.6% | 88,269 | 5.6% | 3,398 | 5.4% | 2,087 | 5.2% |
| $T_{C3}$ | 154 | 10.0% | 154,860 | 9.9% | 5,883 | 9.4% | 3,701 | 9.2% |
| $T_{C4}$ | 67 | 4.4% | 59,598 | 3.8% | 2,434 | 3.9% | 1,525 | 3.8% |
| $T_{C5}$ | 118 | 7.7% | 93,530 | 6.0% | 3,664 | 5.9% | 2,371 | 5.9% |
| $T_{C6}$ | 132 | 8.6% | 127,011 | 8.1% | 4,758 | 7.6% | 2,960 | 7.4% |
| $T_{C7}$ | 176 | 11.4% | 196,128 | 12.5% | 7,943 | 12.7% | 5,055 | 12.6% |
| $T_{C8}$ | 258 | 16.8% | 252,405 | 16.1% | 9,489 | 15.2% | 6,140 | 15.3% |
| $T_{C9}$ | 479 | 31.1% | 546,967 | 35.0% | 22,866 | 36.6% | 15,147 | 37.7% |
| Total | 1,539 |  | 1,564,455 |  | 62,449 |  | 40,202 |  |

**Table S4.** The sum of the proportions for different groups of topologies according to the results of submodel 1 on empirical data C. H - Human; C - Chimpanzee; G - Gorilla; MM - *Macaca mulatta;* MF - *Macaca fascicularis*; MN - *Macaca nemestrina*. Note that the last decimal place of the sums may not match due to the round-up effect.

|  | ((MF,MN),MM) | ((MM,MN),MF) | ((MF,MM),MN) | Sum |
| --- | --- | --- | --- | --- |
| ((G,C),H) | $T_{C1}$<br>0.4% | $T_{C2}$<br>7.0% | $T_{C3}$<br>8.4% | 15.8% |
| ((H,G),C) | $T_{C4}$<br>7.7% | $T_{C5}$<br>2.9% | $T_{C6}$<br>18.3% | 28.9% |
| ((H,C),G) | $T_{C7}$<br>13.0% | $T_{C8}$<br>8.7% | $T_{C9}$<br>33.6% | 55.3% |
| Sum | 21.1% | 18.6% | 60.3% |  |

**Table S5.** The sum of the proportions for different groups of topologies according to the results of submodel 2 on empirical data C. H - Human; C - Chimpanzee; G - Gorilla; MM - *Macaca mulatta;* MF - *Macaca fascicularis;* MN - *Macaca nemestrina.* Note that the last decimal place of the sums may not match due to the round-up effect.

|  | ((MF,MN),MM) | ((MM,MN),MF) | ((MF,MM),MN) | Sum |
| --- | --- | --- | --- | --- |
| ((G,C),H) | $T_{C1}$<br>0.4% | $T_{C2}$<br>10.4% | $T_{C3}$<br>8.2% | 15.8% |
| ((H,G),C) | $T_{C4}$<br>2.1% | $T_{C5}$<br>2.5% | $T_{C6}$<br>14.0% | 28.9% |
| ((H,C),G) | $T_{C7}$<br>13.1% | $T_{C8}$<br>8.4% | $T_{C9}$<br>41.1% | 55.3% |
| Sum | 15.5% | 21.2% | 63.3% |  |

**Table S6.** Detailed information of the genes in empirical dataset D. *T*_*D*1_ is the commonly accepted species tree.

| Topology | Number of genes<br>(Gene tree %) |  | Sum of gene lengths |  | Total number of<br>variable sites |  | Total number of<br>parsimony<br>informative sites |  |
| --- | --- | --- | --- | --- | --- | --- | --- | --- |
| $T_{D1}$ | 295 | 37.06% | 329,864 | 36.06% | 30,823 | 36.72% | 3,064 | 36.69% |
| $T_{D2}$ | 258 | 32.41% | 302,224 | 33.04% | 27,225 | 32.43% | 2,692 | 32.24% |
| $T_{D3}$ | 243 | 30.53% | 282,703 | 30.90% | 25,903 | 30.85% | 2,594 | 31.07% |
| Other | 761 | — | 695,964 | — | 56,689 | — | 5,426 | — |
| $T_{D1} + T_{D2} + T_{D3}$ | 796 | | 914,791 | | 83,951 | | 8,350 | |

## Bibliography

1. Alexandros Stamatakis. RAxML version 8: atoolfor phylogenetic analysis and post-analysis of large phylogenies. Bioinformatics, 30(9):1312–1313, May 2014.

2. Subha Kalyaanamoorthy, Bui Quang Minh, Thomas K F Wong, Arndt von Haeseler, and Lars S Jermiin. ModelFinder: fast model selection for accurate phylogenetic estimates. Nat. Methods, 14(6):587–589, June 2017.

3. Stéphane Guindon, Jean-François Dufayard, Vincent Lefort, Maria Anisimova, Wim Hordijk, and Olivier Gascuel. New algorithms and methods to estimate maximum-likelihood phylogenies: assessing the performance of PhyML 3.0. Syst. Biol., 59(3):307–321, May 2010.

4. Fredrik Ronquist and John P Huelsenbeck. MrBayes 3: Bayesian phylogenetic inference under mixed models. Bioinformatics, 19(12):1572–1574, August 2003.

5. Remco Bouckaert, Timothy G Vaughan, Joëlle Barido-Sottani, Sebastián Duchêne, Mathieu Fourment, Alexandra Gavryushkina, Joseph Heled, Graham Jones, Denise Kühnert, Nicola De Maio, Michael Matschiner, Fábio K Mendes, Nicola F Müller, Huw A Ogilvie, Louis du Plessis, Alex Popinga, Andrew Rambaut, David Rasmussen, Igor Siveroni, Marc A Suchard, Chieh-Hsi Wu, Dong Xie, Chi Zhang, Tanja Stadler, and Alexei J Drummond. BEAST 2.5: An advanced software platform for bayesian evolutionary analysis. PLoS Comput. Biol., 15(4):e1006650, April 2019.

6. Diep Thi Hoang, Le Sy Vinh, Tomáš Flouri, Alexandros Stamatakis, Arndt von Haeseler, and Bui Quang Minh. MPBoot: fast phylogenetic maximum parsimony tree inference and bootstrap approximation. BMC Evol. Biol., 18(1):11, February 2018.

7. Cheng Ye, Bryan Thornlow, Angie Hinrichs, Alexander Kramer, Cade Mirchandani, Devika Torvi, Robert Lanfear, Russell Corbett-Detig, and Yatish Turakhia. matoptimize: A parallel tree optimization method enables online phylogenetics for SARS-CoV-2. Bioinformatics, June 2022.

8. Pablo A Goloboff and Santiago A Catalano. TNT version 1.5, including a full implementation of phylogenetic morphometrics. Cladistics, 32(3):221–238, June 2016.

9. O Gascuel. BIONJ: an improved version of the NJ algorithm based on a simple model of sequence data. Mol. Biol. Evol., 14(7):685–695, July 1997.

10. Vincent Lefort, Richard Desper, and Olivier Gascuel. FastME 2.0: A comprehensive, accurate, and fast Distance-Based phylogeny inference program. Mol. Biol. Evol., 32(10):2798–2800, October 2015.

11. Kevin Howe, Alex Bateman, and Richard Durbin. QuickTree: building huge Neighbour-Joining trees of protein sequences. Bioinformatics, 18(11):1546–1547, November 2002.

12. M Simonsen and C N S Pedersen. Rapid computation of distance estimators from nucleotide and amino acid alignments. In Proceedings of the 2011 ACM Symposium on Applied Computing, SAC ‘11, pages 89–93, New York, NY, USA, March 2011. Association for Computing Machinery.

13. Wayne P Maddison. Gene trees in species trees. Syst. Biol., 46(3):523–536, September 1997.

14. R Nichols. Gene trees and species trees are not the same. Trends Ecol. Evol., 16(7):358–364, July 2001.

15. Scott V Edwards. Is a new and general theory of molecular systematics emerging? Evolution, 63(1):1–19, January 2009.

16. David Bryant, Remco Bouckaert, Joseph Felsenstein, Noah A Rosenberg, and Arindam RoyChoudhury. Inferring species trees directly from biallelic genetic markers: bypassing gene trees in a full coalescent analysis. Mol. Biol. Evol., 29(8):1917–1932, August 2012.

17. Chao Zhang, Maryam Rabiee, Erfan Sayyari, and Siavash Mirarab. ASTRAL-III: polynomial time species tree reconstruction from partially resolved gene trees. BMC Bioinformatics, 19 (Suppl 6):153, May 2018.

18. Liang Liu, Lili Yu, and Scott V Edwards. A maximum pseudo-likelihood approach for estimating species trees under the coalescent model. BMC Evol. Biol., 10:302, October 2010.

19. Julia Chifman and Laura Kubatko. Identifiability of the unrooted species tree topology under the coalescent model with time-reversible substitution processes, site-specific rate variation, and invariable sites. J. Theor. Biol., 374:35–47, June 2015.

20. Joseph Heled and Alexei J Drummond. Bayesian inference of species trees from multilocus data. Mol. Biol. Evol., 27(3):570–580, March 2010.

21. Huw A Ogilvie, Remco R Bouckaert, and Alexei J Drummond. StarBEAST2 brings faster species tree inference and accurate estimates of substitution rates. Mol. Biol. Evol., 34(8):2101–2114, August 2017.

22. Dingqiao Wen, Yun Yu, Jiafan Zhu, and Luay Nakhleh. Inferring phylogenetic networks using PhyloNet. Syst. Biol., 67(4):735–740, July 2018.

23. Claudia Solís-Lemus, Paul Bastide, and Cécile Ané. PhyloNetworks: A package for phylogenetic networks. Mol. Biol. Evol., 34(12):3292–3298, December 2017.

24. Chi Zhang, Huw A Ogilvie, Alexei J Drummond, and Tanja Stadler. Bayesian inference of species networks from multilocus sequence data. Mol. Biol. Evol., 35(2):504–517, February 2018.

25. Tomáš Flouri, Xiyun Jiao, Bruce Rannala, and Ziheng Yang. Species tree inference with BPP using genomic sequences and the multispecies coalescent. Mol. Biol. Evol., 35(10):2585–2593, October 2018.

26. Elizabeth S Allman, John A Rhodes, and Seth Sullivant. When do phylogenetic mixture models mimic other phylogenetic models? Syst. Biol., 61(6):1049–1059, December 2012.

27. Z Yang. Maximum likelihood phylogenetic estimation from DNA sequences with variable rates over sites: approximate methods. J. Mol. Evol., 39(3):306–314, September 1994.

28. Nicolas Lartillot and Hervé Philippe. A bayesian mixture model for across-site heterogeneities in the amino-acid replacement process. Mol. Biol. Evol., 21(6):1095–1109, June 2004.

29. Si Quang Le, Cuong Cao Dang, and Olivier Gascuel. Modeling protein evolution with several amino acid replacement matrices depending on site rates. Mol. Biol. Evol., 29(10):2921–2936, October 2012.

30. Stephen M Crotty, Bui Quang Minh, Nigel G Bean, Barbara R Holland, Jonathan Tuke, Lars S Jermiin, and Arndt Von Haeseler. GHOST: Recovering historical signal from heterotachously evolved sequence alignments, 2019.

31. Bryant and Hahn. The concatenation question. Phylogenetics in the Genomic Era, 2020.

32. Lam-Tung Nguyen, Heiko A Schmidt, Arndt von Haeseler, and Bui Quang Minh. IQ-TREE: a fast and effective stochastic algorithm for estimating maximum-likelihood phylogenies. Mol. Biol. Evol., 32(1):268–274, January 2015.

33. Tavaré. Some probabilistic and statistical problems in the analysis of DNA sequences. Lectures on mathematics in the life sciences, 17:57–86, 1986.

34. J Felsenstein. Evolutionary trees from DNA sequences: a maximum likelihood approach. J. Mol. Evol., 17(6):368–376, 1981.

35. Kenneth P Burnham and David R Anderson. Model Selection and Multimodel Inference: A Practical Information-Theoretic Approach. Springer Science & Business Media, May 2002.

36. A P Dempster, N M Laird, and D B Rubin. Maximum likelihood from incomplete data via theEMAlgorithm. J. R. Stat. Soc., 39(1):1–22, September 1977.

37. R Fletcher. Practical Methods of Optimization. John Wiley & Sons, June 2013.

38. Nhan Ly-Trong, Suha Naser-Khdour, Robert Lanfear, and Bui Quang Minh. AliSim: A fast and versatile phylogenetic sequence simulator for the genomic era. Mol. Biol. Evol., 39(5), May 2022.

39. Richard R Hudson. Generating samples under a Wright–Fisher neutral model of genetic variation. Bioinformatics, 18(2):337–338, February 2002.

40. John Gatesy and Mark S Springer. Phylogenetic analysis at deep timescales: unreliable gene trees, bypassed hidden support, and the coalescence/concatalescence conundrum. Mol. Phylogenet. Evol., 80:231–266, November 2014.

41. Dan Vanderpool, Bui Quang Minh, Robert Lanfear, Daniel Hughes, Shwetha Murali, R Alan Harris, Muthuswamy Raveendran, Donna M Muzny, Mark S Hibbins, Robert J Williamson, Richard A Gibbs, Kim C Worley, Jeffrey Rogers, and Matthew W Hahn. Primate phylogenomics uncovers multiple rapid radiations and ancient interspecific introgression. PLoS Biol., 18(12):e3000954, December 2020.

42. Ingo Ebersberger, Petra Galgoczy, Stefan Taudien, Simone Taenzer, Matthias Platzer, and Arndt von Haeseler. Mapping human genetic ancestry. Mol. Biol. Evol., 24(10):2266–2276, October 2007.

43. Laura Salter Kubatko and James H Degnan. Inconsistency of phylogenetic estimates from concatenated data under coalescence. Syst. Biol., 56(1):17–24, February 2007.

44. Sebastien Roch and Mike Steel. Likelihood-based tree reconstruction on a concatenation of aligned sequence data sets can be statistically inconsistent. Theor. Popul. Biol., 100C: 56–62, March 2015.

45. Fábio K Mendes and Matthew W Hahn. Why concatenation fails near the anomaly zone. Syst. Biol., 67(1):158–169, January 2018.

46. John A Rhodes and Seth Sullivant. Identifiability of large phylogenetic mixture models. Bull. Math. Biol., 74(1):212–231, January 2012.

47. Elizabeth S Allman and John A Rhodes. The identifiability of tree topology for phylogenetic models, including covarion and mixture models. J. Comput. Biol., 13(5):1101–1113, June 2006.

48. Elizabeth S Allman, Sonja Petrović, John A Rhodes, and Seth Sullivant. Identifiability of two-tree mixtures for group-based models. IEEE/ACM Trans. Comput. Biol. Bioinform., 8(3):710–722, May 2011.

49. Joseph Felsenstein. Inferring Phylogenies. Sinauer, 2003.

50. Fábio K Mendes and Matthew W Hahn. Gene tree discordance causes apparent substitution rate variation. Syst. Biol., 65(4):711–721, July 2016.

51. S V Edwards and P Beerli. Perspective: gene divergence, population divergence, and the variance in coalescence time in phylogeographic studies. Evolution, 54(6):1839–1854, December 2000.

52. Richard E Green, Johannes Krause, Adrian W Briggs, Tomislav Maricic, Udo Stenzel, Martin Kircher, Nick Patterson, Heng Li, Weiwei Zhai, Markus Hsi-Yang Fritz, Nancy F Hansen, Eric Y Durand, Anna-Sapfo Malaspinas, Jeffrey D Jensen, Tomas Marques-Bonet, Can Alkan, Kay Prüfer, Matthias Meyer, Hernán A Burbano, Jeffrey M Good, Rigo Schultz, Ayinuer Aximu-Petri, Anne Butthof, Barbara Höber, Barbara Höffner, Madlen Siegemund, Antje Weihmann, Chad Nusbaum, Eric S Lander, Carsten Russ, Nathaniel Novod, Jason Affourtit, Michael Egholm, Christine Verna, Pavao Rudan, Dejana Brajkovic, Željko Kucan, Ivan Gušic, Vladimir B Doronichev, Liubov V Golovanova, Carles Lalueza-Fox, Marco de la Rasilla, Javier Fortea, Antonio Rosas, Ralf W Schmitz, Philip L F Johnson, Evan E Eichler, Daniel Falush, Ewan Birney, James C Mullikin, Montgomery Slatkin, Rasmus Nielsen, Janet Kelso, Michael Lachmann, David Reich, and Svante Pääbo. A draft sequence of the neandertal genome. Science, 328(5979):710–722, May 2010.

53. Asger Hobolth, Ole F Christensen, Thomas Mailund, and Mikkel H Schierup. Genomic relationships and speciation times of human, chimpanzee, and gorilla inferred from a coalescent hidden markov model. PLoS Genet., 3(2):e7, February 2007.

54. D J Balding, R A Nichols, and D M Hunt. Detecting gene conversion: primate visual pigment genes. Proc. Biol. Sci., 249(1326):275–280, September 1992.

